# *HASTER* is a transcriptional stabilizer of *HNF1A*

**DOI:** 10.1101/2021.05.12.443907

**Authors:** Anthony Beucher, Irene Miguel-Escalada, Diego Balboa, Matías G. De Vas, Miguel Angel Maestro, Javier Garcia-Hurtado, Aina Bernal, Roser Gonzalez-Franco, Pierfrancesco Vargiu, Holger Heyn, Philippe Ravassard, Sagrario Ortega, Jorge Ferrer

**Affiliations:** Section of Genetics and Genomics, Department of Metabolism, Digestion and Reproduction, Imperial College London, London W12 0NN, UK; Centre for Genomic Regulation (CRG), The Barcelona Institute of Science and Technology (BIST), 08003 Barcelona, Spain; Centro de Investigación Biomédica en Red de Diabetes y Enfermedades Metabólicas Asociadas (CIBERDEM), Spain; Transgenics Unit, Spanish National Cancer Research Centre-CNIO, 28029 Madrid, Spain; CNAG-CRG, Centre for Genomic Regulation (CRG), Barcelona Institute of Science and Technology (BIST), Barcelona, Spain; Institut du Cerveau et de la Moelle, Biotechnology & Biotherapy team, CNRS UMR7225; INSERM U975; University Pierre et Marie Curie, 75005 Paris, France

## Abstract

The biological purpose and disease relevance of long noncoding RNAs (lncRNAs) is poorly understood. We examined *HASTER*, a lncRNA antisense to *HNF1A*. Haploinsufficient mutations in *HNF1A*, encoding a homeodomain transcription factor, cause diabetes mellitus. Using mouse and human models, we show that *HASTER* maintains HNF1A at cell-specific physiological concentrations through positive and negative feedback loops. *Haster* mutant pancreatic β cells thus showed variegated HNF1A overexpression or silencing, causing insulin-deficiency and diabetes. We demonstrate that the *HASTER* promoter acts in *cis* to prevent *HNF1A* overexpression and silencing, and link *HASTER*-dependent inhibition to local remodelling of 3D chromatin architecture. We further show that *HASTER* negative feedback ensures that HNF1A creates open chromatin at appropriate cell-specific genome regions. Our studies expose a *cis*-regulatory element that is unlike enhancers or silencers, and instead stabilizes expression levels of a pioneer transcription factor. They also show that disruption of a mammalian lncRNA can cause diabetes mellitus.

---

The transcription of genes is controlled by *cis*-acting promoter and enhancer sequences, many of which harbor disease variants. Mammalian genomes also contain >20,000 long non-coding RNAs (lncRNAs)^1,2^. Although the function of most lncRNAs has not been explored, some lncRNAs are known to regulate gene transcription^3,4^. A significant fraction of lncRNAs is transcribed from evolutionary conserved promoters located near genes encoding lineage-specific regulators^3,5–7^, suggesting that they can function as *cis*-regulators. The significance of these observations is, however, uncertain. For several lncRNAs, knockdown experiments have revealed transcriptional effects on nearby genes^8–10^, while in-depth genetic analysis of selected loci has demonstrated *bona fide cis*-regulatory functions of lncRNAs^3,11–16^. There are nevertheless still major gaps in our understanding of the regulatory purpose of lncRNAs, and how they are fundamentally different from more established gene regulatory elements. Furthermore, the extent to which genetic disruption of *cis*-regulatory lncRNAs can lead to physiologically relevant phenotypes is unclear. Such questions are central to dissect the possible role of lncRNAs in differentiation and human disease.

In this study we have examined *HASTER*, a lncRNA at the *HNF1A* locus. Mutations in *HNF1A*, encoding a homeodomain transcription factor^17^, cause MODY3, the most frequent form of monogenic diabetes mellitus^18^, while rare and common variants predispose to type 2 diabetes^19,20^. Homozygous *Hnf1a* null mutant mice have shown that HNF1A is essential for differentiation programs in various organs, whereas human *HNF1A* haploinsufficiency causes diabetes due to selective abnormalities in pancreatic β cells, indicating that *HNF1A* exhibits cell-specific gene dosage sensitivity^18,21–26^. Our study now shows that *HASTER* is a cell-specific transcriptional stabilizer of HNF1A concentrations. We demonstrate that stabilization is achieved through a dual mechanism in which the *HASTER* promoter prevents silencing of the *HNF1A* locus, but also acts as a negative feedback to prevent HNF1A overexpression and binding to inappropriate genomic regions. We further show that disruption of this stabilizing function causes diabetes mellitus. These findings warrant a need to interrogate *cis*-acting lncRNA sequences to decipher the genetic underpinnings of diabetes mellitus.

## Results

### Evolutionary conserved co-expression of *HNF1A* and *HASTER*

We focused on a lncRNA that is transcribed antisense to *HNF1A* (also known as *HNF1A-AS1*, or *Hnf1a-os1* and *2* in mouse) (**Fig. 1a**). We refer to *HNF1A* antisense transcripts and their promoter DNA as *HASTER*, for *<u>H</u>NF1<u>A</u>* <u>st</u>abiliz<u>er</u>. *HASTER* transcripts have been previously proposed to exert *trans*-regulation of proliferation in cell-based models^14,27–31^, but so far the transcriptional regulatory function and sequence determinants of *HASTER* have not been characterized with genetic tools. *HASTER* has numerous transcript isoforms that originate from a major upstream transcriptional start site in human islets, and an additional downstream start site in other tissues (**Supplementary Fig. 1a,b**). Both transcriptional start sites are located in adjacent evolutionary conserved sequences that show active promoter chromatin in islets and liver, with very high H3K4me3 and low H3K4me1 in downstream nucleosomes (**Supplementary Fig. 1b**).

**Fig. 1.**
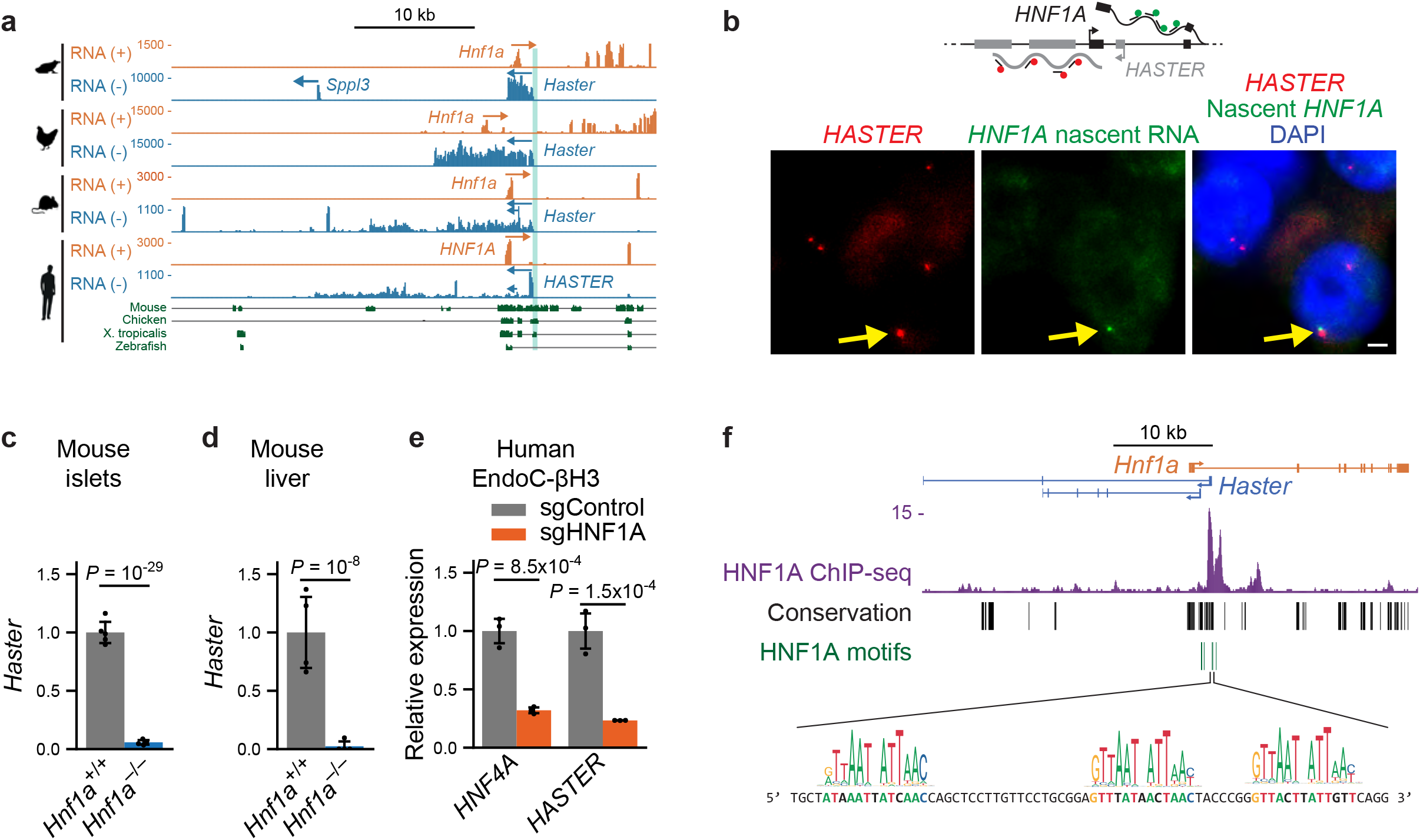
*HASTER* is an evolutionary conserved nuclear RNA regulated by *HNF1A*. **a**, Liver strand-specific RNA-seq and Multiz Alignments in the indicated species. **b**, Single molecule fluorescent in situ hybridization for *HASTER* (exonic probes) and *HNF1A* nascent transcripts (intronic probes) in EndoC-βH3 β cells. Yellow arrows indicate co-localization of *HASTER* and nascent *HNF1A* transcripts. Scale bar, 20 μm. **c**,**d**, *Haster* was decreased in *Hnf1a*^−/−^ islets (n = 4 *Hnf1a*^−/−^ mice and n = 5 *Hnf1a*^+/+^ mice) and liver (n = 4 mice per genotype); mean ± s.d., Wald test adjusted *P*. **e**, EndoC-βH3 cells carrying indels in *HNF1A* first exon showed decreased *HASTER* and *HNF4A*, an HNF1A-dependent gene; n=3 lentiviral transductions, mean ± s.d., normalized by *TBP*. *P*-values are from two-tailed Student’s t-test. **f**, HNF1A binds the *Haster* promoter in mouse liver. Major start sites in GENCODE *Haster* transcripts are shown (see also **Supplementary Fig. 1**), as well as seven HNF1A motifs (JASPAR CORE > 0.8). Sequences of three motifs are highlighted below.

*HASTER* is expressed exclusively in *HNF1A*-expressing tissues, including liver, gut, pancreas and kidney, and has the same antisense configuration across species (**Fig. 1a, Supplementary Fig. 1c,d)**. Subcellular fractionation of EndoC-βH3 human β cells showed that *HASTER* transcripts were associated with chromatin, and single molecule fluorescence in situ hybridization showed that *HASTER* transcripts were exclusively present in one or two nuclear foci that co-localized with *HNF1A* nascent transcripts (**Fig. 1b, Supplementary Fig. 2**). *HASTER*, therefore, is an evolutionary conserved nuclear lncRNA that is co-expressed with *HNF1A* across tissues. These findings suggest a regulatory interaction between *HASTER* and *HNF1A*.

### HNF1A is a positive regulator of *HASTER*

To explore functional relationships between *HNF1A* and *HASTER*, we first examined *HASTER* expression in *HNF1A*-deficient cells. *HASTER* was strongly downregulated in pancreatic islets and liver from *Hnf1a*^−/−^ mice that carried a germline indel in the *Hnf1a* coding sequence, as well as in *HNF1A*-deficient EndoC-βH3 human β cells (**Fig. 1c-e**). *HASTER* lncRNA seemed highly sensitive to *HNF1A* expression, because partial *HNF1A* knockdown caused markedly decreased *HASTER* and only marginal changes in other HNF1A-dependent genes like *HNF4A*^32^(**Supplementary Fig. 3a**). Conversely, upregulation of *Hnf1a* mRNA by ~30-80% through CRISPR/Cas9 Synergistic Activation Mediator (CRISPR-SAM) led to ~50-120% increased *Haster* RNA (**Supplementary Fig. 3b**). This effect was likely direct, because the *HASTER* promoter has seven HNF1A recognition sequences that are strongly bound by HNF1A in mouse liver and EndoC-βH3 human β cells (**Fig. 1f**). These findings suggested that the *HASTER* promoter could function as an HNF1Asensing platform that drives *HASTER* RNA transcription in accordance with the cellular concentration of HNF1A.

### *HASTER* is a negative regulator of *HNF1A*

We next examined whether *HASTER* in turn influences *HNF1A* expression. We created a 320 bp deletion of the main *HASTER* promoter (P1) in human Embryonic Stem Cells (hESCs)(**Fig. 2a**), and generated hepatocyte-like cells using established differentiation protocols^33^. *HASTER* was detected at maximal levels in control hepatoblasts, while *HNF1A* was detected at low levels and then increased upon maturation to hepatocytes (**Fig. 2b**). *HASTER* mutant cells expectedly showed near-complete elimination of *HASTER*, and this was accompanied by increased hepatocyte *HNF1A* mRNA (mean 1.3 and 1.6-fold vs. control cells for independent deletions, *P* = 0.01 and 0.04, Student’s t-test) (**Fig. 2b**). This suggested that *HASTER* exerts negative regulation of *HNF1A* in an *in vitro* human liver cell model.

**Fig. 2.**
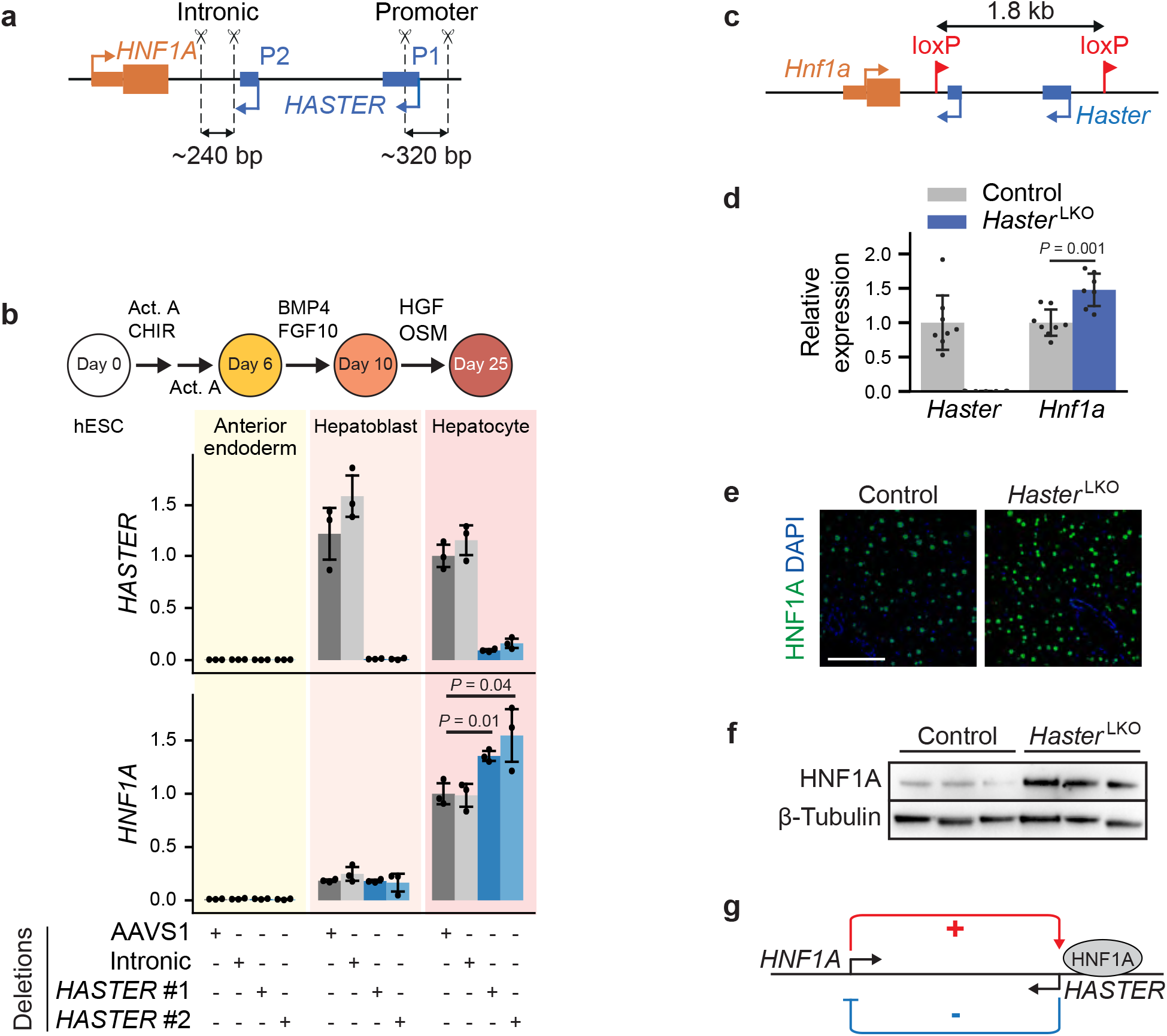
*HASTER* is a negative regulator of *HNF1A* in mouse and human hepatocytes. **a**, Homozygous deletions of the *HASTER* promoter (two deletions with independent sgRNA pairs) or control deletions in *HNF1A* intron 1 or AAVS1 were generated in hESCs. **b**, *HNF1A* mRNA was increased in differentiated hepatocytes from *HASTER* mutant hESCs. n = 3 independent clones per deletion. Bar graphs show *RPLP0*-normalized mean ± s.d., two-tailed Student’s t-test. **c**, Schematic of the mouse *Haster* ^f^ allele. **d**, Liver RNA levels in 7 *Haster* ^LKO^ and 8 control mice. *Tbp*-normalized mean ± s.d., two-tailed Student’s t-test. **e**, Liver HNF1A immunofluorescence in indicated genotypes. **f**, Western blot for HNF1A on liver extracts. N = 3 mice for each genotype. **g**, Schematic of the *HNF1A/HASTER* negative feedback loop.

To examine this regulatory interaction *in vivo*, we generated mice with LoxP sites flanking a 1.8 Kb region containing both *Haster* transcriptional start sites (**Fig. 2c, Supplementary Fig. 4**), and crossed them with a liver-expressing Cre transgenic^34^ to generate homozygous liver-specific *Haster* promoter deletions (*Haster*^LKO^). *Haster*^LKO^ mice were born at Mendelian rates, showed normal organ formation, weight, and glucose homeostasis (**Supplementary Fig. 5a,b**). Consistent with the hESC differentiation model, *Haster*^LKO^ mice showed increased liver *Hnf1a* mRNA and protein (**Fig. 2d-f**). Similar results were observed in liver from homozygous germline *Haster* mutant mice (**Supplementary Fig. 5c**). Taken together, our observations showed that HNF1A positively regulates *HASTER*, while *HASTER* negatively regulates *HNF1A* forming a negative feedback loop that is conserved in mouse and human cells (**Fig. 2g**).

### *HASTER* negative feedback controls HNF1A pioneer activity

To investigate the consequences of disrupting the *HNF1A-HASTER* negative feedback, we performed RNA-seq on liver from *Haster*^LKO^ and control mice. This revealed profound transcriptional changes in *Haster*^LKO^ liver (**Fig. 3a and Supplementary Table 1**). Consistent with increased HNF1A levels in *Haster*^LKO^ liver, deregulated transcripts and functional annotations were negatively correlated with those of *Hnf1a* knockout liver^22^(**Fig. 3b, Supplementary Fig. 5d)**. A subset of genes that were most strongly upregulated in *Haster*^LKO^ liver were, however, specifically expressed in kidney or intestine, two organs that also express HNF1A at high levels (**Fig. 3c**, **Supplementary Fig. 6a**). *Haster* mutations, therefore, led to increased expression of HNF1A-dependent liver genes, but also activated an ectopic transcriptional program.

**Fig. 3.**
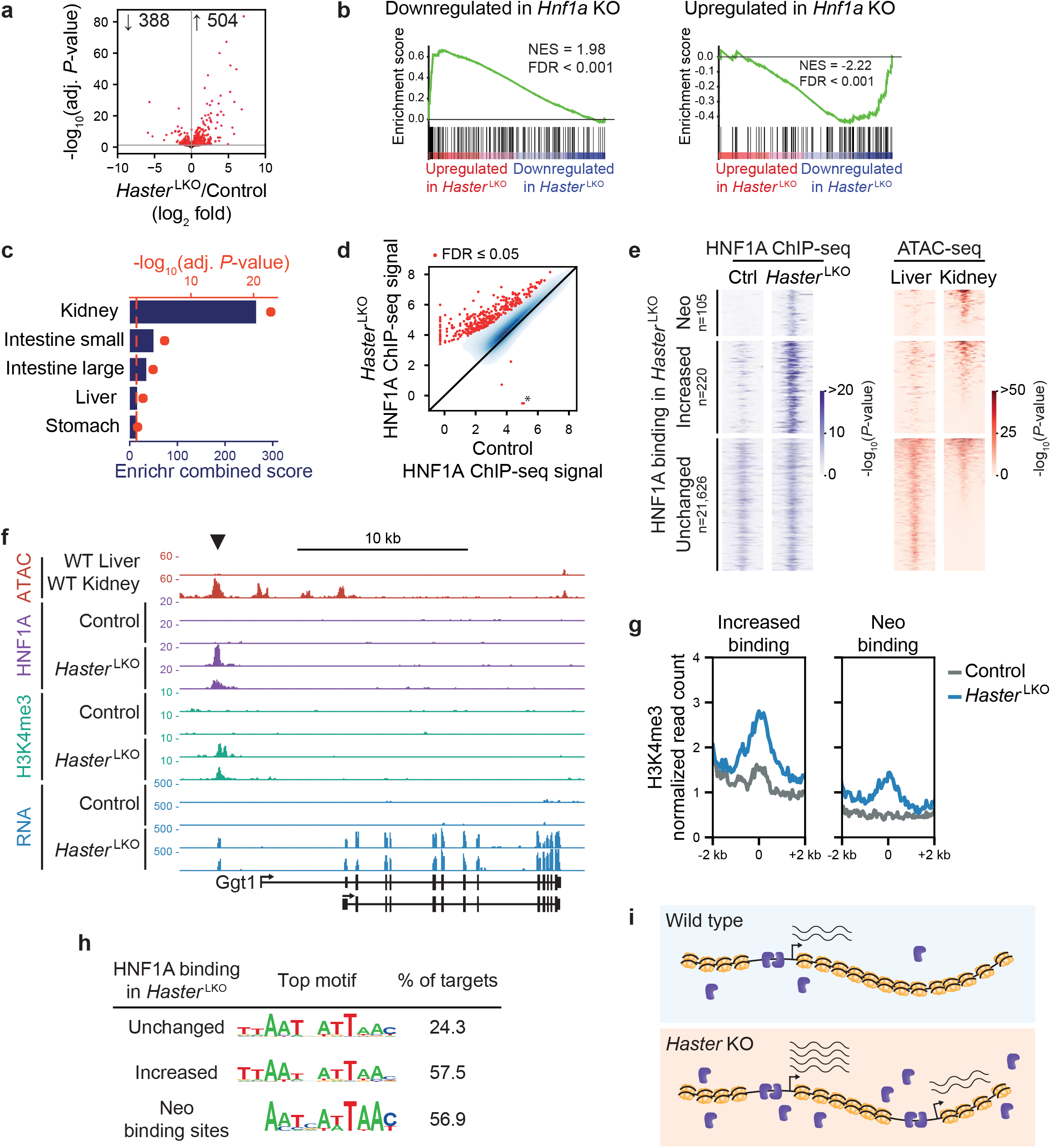
*Haster* controls HNF1A pioneer activity. **a**, RNA-seq in *Haster* ^LKO^ liver. Differentially expressed genes (adjusted *P* ≤ 0.05) are highlighted in red, and total numbers are indicated; n = 5 mice per genotype. **b**, GSEA showing that genes up- or downregulated in *Hnf1a* KO liver have opposite expression patterns in *Haster* ^LKO^ liver. **c**, Enrichment of *Haster* ^LKO^ liver upregulated genes in different mouse tissues (Mouse Gene Atlas). Bars indicate Enrichr scores and red dots are −log_10_ adjusted *P*-values. **d**, HNF1A binding strength (log_2_ ChIP-seq normalized read count) in *Haster* and control liver (n = 3 mice). Red, differentially bound sites (FDR ≤ LKO 0.05); blue, kernel density of HNF1A bound sites with FDR > 0.05. The asterisk denotes the *Haster* ^LKO^ deletion. **e**, HNF1A occupancy in control and *Haster* ^LKO^ liver (left), and chromatin accessibility for the same regions in liver and kidney (right). Neo binding sites are bound by HNF1A only in *Haster* ^LKO^. Increased bound sites are all the other sites showing increased binding in *Haster* ^LKO^. Heatmaps show average signal of 3 replicates for ChIP-seq and 2 replicates for ATAC-seq, peak centers ± 1kb. **f**, Activation of a kidney-specific gene in *Haster* ^LKO^ liver. **g**, H3K4me3 in HNF1A-bound regions in *Haster* ^LKO^ and controls (average of three mice). **h**, Top HOMER de-novo motif for the different categories of HNF1A peaks. **i**, Model showing that *Haster* KO leads to increased HNF1A (blue), causing increased HNF1A binding and expression of HNF1A-bound genes (bottom left), and HNF1A neo-binding sites at inaccessible chromatin that activate an ectopic gene program (bottom right).

We next examined HNF1A genomic binding in *Haster*^LKO^ liver. Overall, HNF1A binding strength was increased in *Haster*^LKO^ liver; 325 peaks showed increased HNF1A binding at *FDR* ≤ 0.05, whereas only 2 peaks showed decreased binding (**Fig. 3d**). Importantly, *Haster*^LKO^ liver showed 105 HNF1A binding sites in regions that were not bound by HNF1A in control livers, hereafter referred to as HNF1A neo-binding sites (**Fig. 3d-f)**.

HNF1A can bind *in vitro* to nucleosomal DNA^35^, and has been used to reprogram fibroblasts into hepatocytes^36^, which are properties of pioneer transcription factors^37^. Although pioneer transcription factors can bind inaccessible chromatin, they typically bind different genomic regions across tissues^22,38^, suggesting that cell-specific parameters, such as perhaps cellular transcription factor concentrations, might influence their *in vivo* binding selectivity. In keeping with this notion, HNF1A neo-binding sites did not show accessible chromatin in normal liver (**Fig. 3e,f**), whereas they showed classical active chromatin modifications (H3K4me3 and H3K27ac) in *Haster*^LKO^ liver (**Fig. 3g, Supplementary Fig. 6b-f)**. Interestingly, HNF1A neo-binding sites contained canonical high-affinity HNF1 binding motifs, suggesting that many could be *bona fide* HNF1A targets in other HNF1A-expressing tissues (**Fig. 3h**). Thus, increased HNF1A in *Haster*^LKO^ liver resulted in the creation of new binding sites, which led to the formation of new active chromatin regions.

Increased HNF1A binding at pre-existing active promoters led to increased mRNA in *Haster*^LKO^ liver; ~1/4 of genes in this class showed >2-fold higher expression in *Haster*^LKO^ (**Supplementary Fig. 6d)**. HNF1A neo-binding events in newly generated promoter regions often led to ectopic activation of genes that are normally not expressed in liver, such as kidney-enriched genes *Ggt* and *Tinag* (**Fig. 3f, Supplementary Fig. 6d,e**). Consistently, several HNF1A neo-binding sites did not show accessible chromatin in normal liver yet showed accessible chromatin in other HNF1A-expressing tissues such as kidney (**Fig. 3c,f, Supplementary Fig**. **6a,e**). Some newly activated promoters did not overlap with any annotated mouse transcription start site, suggesting that increased HNF1A could also activate aberrant *de novo* promoters (**Supplementary Fig. 6f,g**).

In summary, genetic disruption of the *HASTER* feedback loop led to increased cellular HNF1A concentrations, which caused either super-activation of pre-existing HNF1A-bound promoters, or the transformation of silent inaccessible chromatin into active promoters (**Fig. 3i**). This indicates that the *HASTER* feedback is crucial to control the pioneering activity of HNF1A, and to fine-tune the tissue specificity of HNF1A-dependent transcriptional programs.

### *Haster* inactivation causes diabetes

Human *HNF1A* haploinsufficiency leads to pancreatic β cell dysfunction and diabetes^18^. To examine the function of *Haster* in pancreatic cells, we used a *Pdx1-* Cre transgene that excised *Haster* in all pancreatic epithelial lineages, and refer to these as *Haster*^pKO^ mice. *Haster*^pKO^ mice showed normal growth and no gross morphological abnormalities **(Supplementary Fig. 7a**), yet displayed glucose intolerance with marked insulin-deficiency by 8 weeks, as well as fasting hyperglycemia (glycemia 137 ± 16 mM in *Haster*^pKO^, 87 ± 5 mM in *Haster*^f/f^ littermates, 98 ± 4 mM in *Pdx1*-Cre; t-test P < 0.05) (**Fig. 4a**,**b, Supplementary Fig. 7b**). Mice with germline mutations (*Haster*^−/−^) were born at Mendelian rates and showed no overt manifestations, but also showed diabetes, glucose intolerance, and hypoinsulinemia at 8 weeks of age (**Fig. 4c-e, Supplementary Fig. 7c,d**). Thus, inactivation of *Haster* in the pancreas or germline led to impaired insulin secretion and diabetes.

**Fig. 4.**
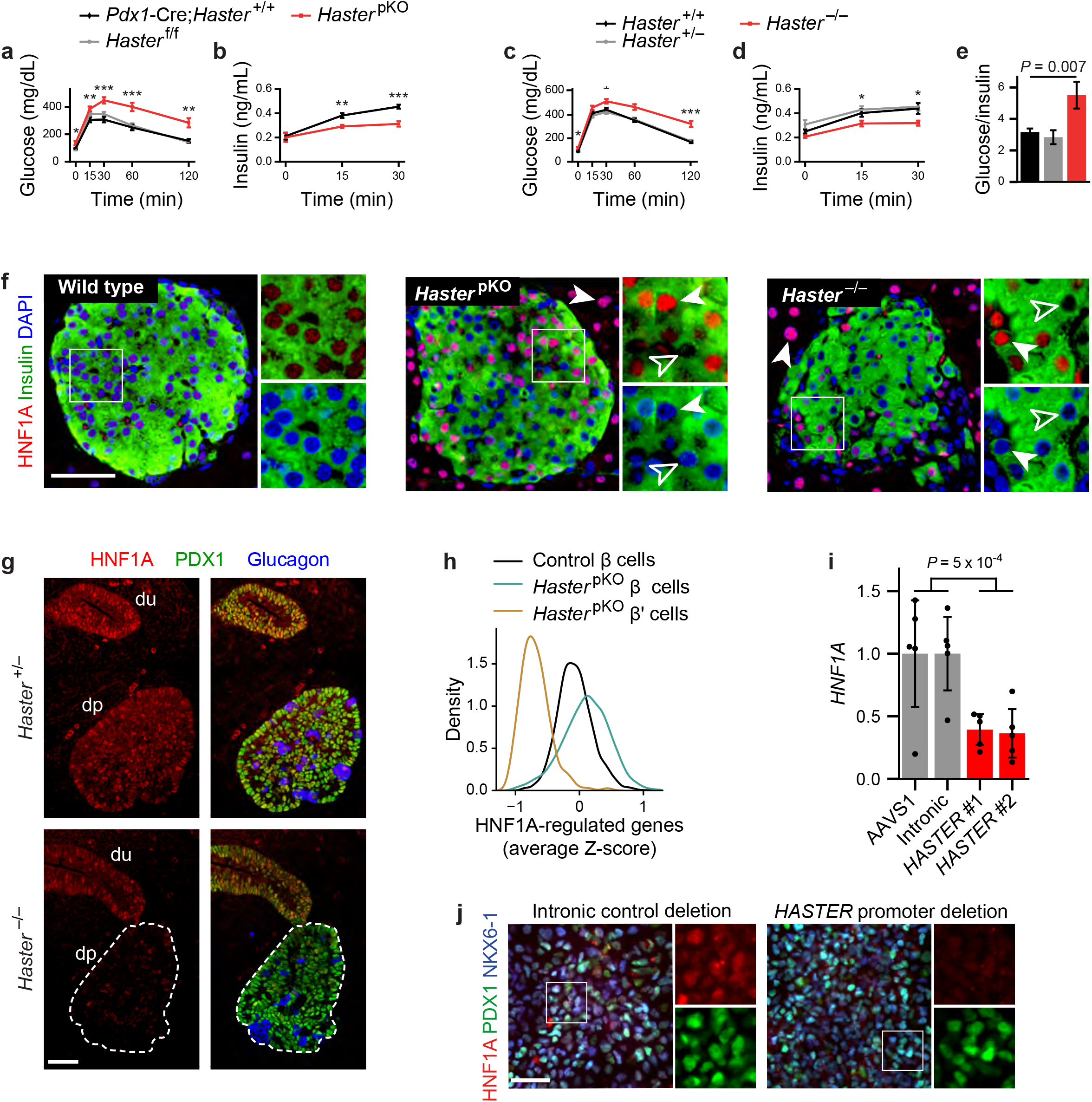
*Haster* deletion causes diabetes through islet-cell HNF1A hyperactivation or silencing. **a**, Intraperitoneal glucose tolerance in 8-week-old male mice (n = 8 *Haster* ^pKO^, n = 12 *Pdx1*-Cre;*Haster* ^+/+^, n = 8 *Haster* ^f/f^). **b**, Plasma insulin of 8-week-old male mice (n = 7 *Haster* ^pKO^, n = 6 *Pdx1*-Cre;*Haster* ^+/+^). **c**, Intraperitoneal glucose tolerance in 8-week-old male mice (n = 9 *Haster* ^−/−^, n = 12 *Haster* ^+/−^, n = 13 *Haster* ^+/+^). **d**, Plasma insulin in 8-week-old male mice (n = 7 *Haster* ^−/−^, n = 6 *Haster* ^+/+^, n = 6 *Haster* ^+/−^). **e**, Glucose-to-insulin ratio in 8-week-old male mice (n = 9 *Haster* ^−/−^, n = 12 *Haster* ^+/−^, n = 13 *Haster* ^+/+^). **a-e**, Mean ± s.e.m., \**P* ≤ 0.05, \*\**P* ≤ 0.01, \*\*\**P* ≤ 0.001. Two-tailed Student’s t-test. **f**, Immunofluorescence for HNF1A and insulin in *Haster* ^+/+^, *Haster* ^pKO^, and *Haster* ^−/−^ adult knockouts, showing either HNF1A overexpression (solid arrows), or no HNF1A expression (hollow arrows) in all endocrine cells of *Haster* ^pKO^ and *Haster* ^−/−^. All acinar cells from *Haster* ^pKO^ and *Haster* ^−/−^ mice overexpressed HNF1A. **g**, Immunofluorescence for HNF1A, the pancreatic and duodenal marker PDX1, and glucagon in *Haster* ^−/−^ and control E11.5 embryos, showing low and heterogeneous HNF1A expression in pancreatic but not gut progenitors. du, duodenum; dp dorsal pancreas. **h**, Kernel density estimation of HNF1A-regulated gene expression (average Z-score) showing either down or upregulation of HNF1A-regulated genes in two *Haster* ^pKO^ β cell clusters (β’ and β, respectively). **i**, *HNF1A* mRNA in hESC-derived pancreatic progenitors carrying *HASTER* P1 homozygous deletions (see **Fig. 2a**). n = 5 independent differentiations. *TBP*-normalized mean ± s.d., two-tailed Student’s t-test. **j**, Immunofluorescence for HNF1A and pancreatic progenitor markers PDX1 and NKX6-1 in hESC-derived pancreatic progenitors carrying the indicated deletions, showing downregulation of HNF1A. **f**,**g**, Scale bar, 50 μm.

### *Haster* KO leads to *HNF1A* hyperactivation or silencing in islet cells

Immunofluorescence analysis of *Haster*^pKO^ and *Haster*^−/−^ pancreas showed that HNF1A expression was strongly upregulated in all acinar cells, and in many endocrine cells (**Fig. 4f**). This confirmed that *Haster* also acts as a negative regulator of *Hnf1a* in the pancreas. However, we also observed that numerous islet endocrine cells from 8- to 12-week-old *Haster*^pKO^ and *Haster*^−/−^ mice were completely devoid of HNF1A immunoreactivity (**Fig. 4f**).

To further understand *Haster*-dependent regulation of pancreatic HNF1A expression, we analyzed embryonic or adult mice in which *Haster* was deleted in the germline (*Haster*^−/−^), in early pancreatic progenitors (*Haster*^pKO^), or in insulin-expressing β cells (*Haster*^βKO^). In embryonic stage E11.5, most *Haster*^−/−^ multipotent pancreatic progenitors showed markedly heterogeneous HNF1A expression, with many cells showing low or no HNF1A expression, whereas HNF1A expression was uniform in surrounding primitive gut cells (**Fig. 4g**). At embryonic stage E15.5, β-cells from *Haster*^−/−^ and *Haster*^pKO^ embryos also showed highly variable HNF1A levels, ranging from an apparent absence to marked overexpression in 1-5% of β cells (**Supplementary Fig. 8a-d**). This contrasted with highly uniform HNF1A staining in control embryonic β cells (**Supplementary Fig. 8a,b**). This dual phenotype became more evident in adult *Haster*^pKO^ and *Haster*^−/−^ mice, which showed more HNF1A-negative cells (62% and 80%, respectively), and more HNF1A overexpressing cells (24% and 10%, respectively) (**Fig. 4f**). Inactivation of *Haster* after the formation of β cells, however, resulted in fewer HNF1A-negative β cells and more frequent HNF1A overexpression (**Supplementary Fig. 8a,f-g**). Thus, *Haster* inactivation caused a unique variegated HNF1A expression phenotype in β cells, with co-existing silencing and overexpression. This unexpected pancreatic endocrine phenotype showed that *Haster* acts as a negative regulator of HNF1A in the pancreas, as in the liver, but further uncovered a developmental cell-specific role of *Haster* to ensure HNF1A expression in early pancreatic progenitors and islet cells. It also revealed that *Haster* is essential for β cell function and glucose homeostasis.

### Single-cell genomics reveals variegated *Haster*-deficient transcriptomes

We next defined the transcriptional impact of HNF1A expression heterogeneity. We performed single-cell RNA-seq of islet cells from three *Haster*^pKO^ and control mice, and used graph-based clustering to separate major endocrine cell types (**Supplementary Fig. 9a-d**). For each cell, we then calculated the average normalized expression of known HNF1A-regulated genes. Consistent with HNF1A expression heterogeneity in *Haster*^pKO^ β cells, we observed increased variability of HNF1A-regulated genes across *Haster*^pKO^ β cells (0.53 and 0.34 interquartile ranges for *Haster*^pKO^ and control β cells, respectively, Brown–Forsythe *P* < 10^−93^)(**Fig. 4h, Supplementary Fig. 9e-g**). Further examination revealed a large cluster of *Haster*^pKO^ β cells with increased expression of HNF1A-regulated genes, and another β cell cluster (β’) with strong downregulation of HNF1A-regulated genes, including well known HNF1A targets such as *Ttr*, *Tmem27*, *Slc2a2*, and *Kif12* (**Fig. 4h, Supplementary Fig. 10 and Supplementary Table 2,3**). Thus, *Haster* deletion *in vivo* caused either functional *HNF1A* deficiency in pancreatic β cells, which is known to cause diabetes, or overexpression of *HNF1A*-dependent genes. *Haster*, therefore, acts to ensure the stability of β cell HNF1A-regulated programs.

### *HASTER* modulates *HNF1A* expression levels in human pancreatic progenitors

We next investigated whether *HASTER* also regulates *HNF1A* in human pancreatic cells. Analysis of published datasets showed that *HASTER* is activated during earliest stages of hESC-derived pancreatic differentiation^39^(**Supplementary Fig. 11a)**. To test HASTER function in human pancreatic progenitors, we used the hESC clones carrying a 320 bp *HASTER* P1 promoter mutation described in **Fig. 2a**, and generated pancreatic progenitors *in vitro^40^*. In contrast with results after hepatic differentiation, which showed increased *HNF1A* mRNA, *HASTER* knock-out pancreatic progenitors showed a 62% decrease of *HNF1A* mRNA, and low heterogenous HNF1A protein levels (**Fig. 4i,j, Supplementary Fig. 11b)**. These results demonstrated that *HASTER* also acts as an organ-specific positive modulator of *HNF1A* in human early pancreatic multipotent progenitor cells.

### The *HASTER* promoter activates *HNF1A* in *cis*

We next explored how *HASTER* exerts positive and negative regulation of *HNF1A*, and first focused on the positive regulatory function. To assess if *HASTER* acts in *cis* or *trans*, we bred compound heterozygous *Hnf1a*^+/−^;*Haster*^+/ −^ mice. Single heterozygous *Haster*^+/−^ or *Hnf1a*^+/−^ mice do not develop hyperglycemia^21^ (in contrast to human *HNF1A* heterozygous mutations, which cause diabetes) (**Fig. 5a**). Remarkably, compound heterozygous *Hnf1a*^+/−^;*Haster*^+/−^ young mice developed severe fasting and fed hyperglycemia with hypoinsulinemia, but otherwise did not exhibit typical manifestations of germ-line *Hnf1a* knock our mice (**Fig. 5a**). This was accompanied by absent HNF1A expression in most β cells of 10 week-old *Hnf1a*^+/−^;*Haster*^+/ −^ mice (**Fig. 5b**), consistent with a lack of HNF1A expression from the *Hnf1a* null allele, and a requirement for *Haster* to maintain wildtype *Hnf1a* in *cis* in islet cells. We also created hybrid strain mice with a heterozygous *Haster* null allele, and found decreased islet *Hnf1a* mRNA in chromosomes carrying the *Haster* null allele (P < 0.02) (**Fig. 5c**). Genetic experiments thus showed that *Haster* acts in *cis* to maintain *Hnf1a* expression in islet β cells.

**Fig. 5.**
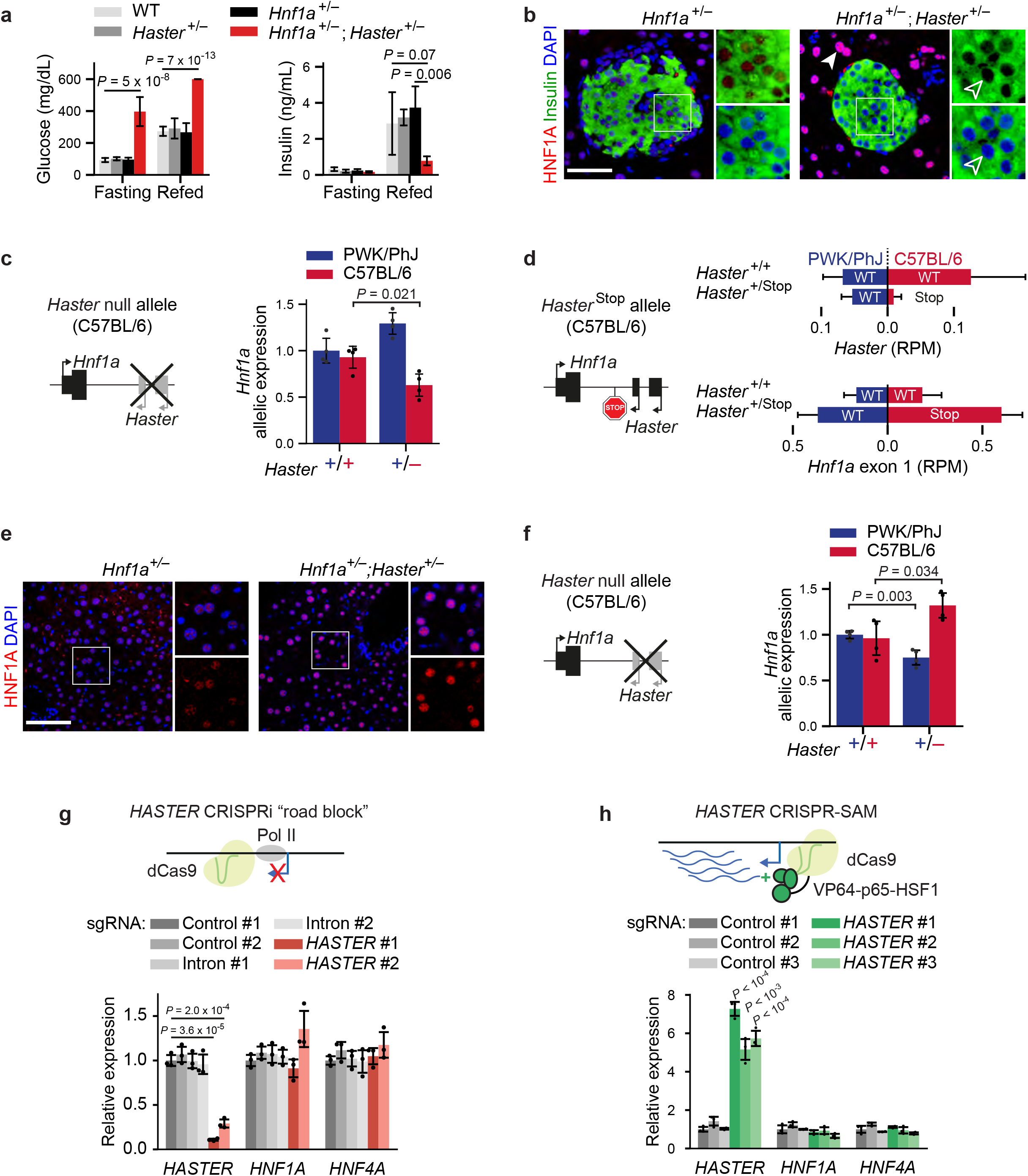
The *HASTER* promoter is a positive and negative *cis*-acting element. **a**, Severe fasting and fed hyperglycemia (left panel; n = 10 to 13 mice per genotype), and reduced insulin secretion (right panel; n = 5 mice per genotype), in *Hnf1a*^+/−^;*Haster* ^+/−^ compound heterozygotes. Mean ± s.d., two-tailed Student’s t-test. **b**, Immunofluorescence shows normal HNF1A in *Hnf1a*^+/−^ islets and no expression in most islet cells from adult *Hnf1a*^+/−^;*Haster* ^+/−^ mice. Scale bar, 50 μm. **c**, Allele-specific *Hnf1a* mRNA in islets from hybrid strain mice that carry the *Haster* mutation in the C57BL/6 chromosome. *Hnf1a* was quantified with strain-specific qPCR and normalized by *Tbp*. n = 4 mice per genotype; mean ± s.d.; two-tailed Student’s t-test, **d**, Strain-specific RNA-seq analysis from *Haster* ^+/Stop^ and *Haster* ^+/+^ PWK/PhJ;C57BL/6 hybrid islets (n=4 mice per genotype). **e**, HNF1A overexpression in liver from *Hnf1a*^+/−^;*Haster* ^+/−^ mice. **f**, Allele-specific *Hnf1a* mRNA in liver from *Haster* ^+/−^ hybrid strain mice that carry the *Haster* mutation in the C57BL/6 chromosome. *Hnf1a* was quantified with strain-specific assays, and normalized by *Tbp*. n = 4 mice per genotype; mean ± s.d.; two-tailed Student’s t-test. **g**, Targeting dCAS9 to *HASTER* transcriptional start site blocked *HASTER* transcription in EndoC-βH3 cells but did not affect *HNF1A* or *HNF4A* mRNAs (n=3 lentiviral transductions). **h**, CRISPR-SAM *HASTER* activation in EndoC-βH3 cells did not affect *HNF1A* and *HNF4A* (n = 3 lentiviral transductions).

We next examined whether *HASTER* transcriptional elongation, RNA molecules, or its promoter DNA are required for the *cis*-regulatory function that prevents *HNF1A* silencing. To this end, we created an allele with a transcriptional termination signal downstream of *Haster*, without altering the *Haster* promoter sequence (*Haster*^Stop^; **Fig. 5d**). We then bred this *Haster*^Stop^ allele in a hybrid strain background, and performed RNA-seq for strain-specific quantitation of *Hnf1a* mRNA in islets. As expected, we found severely diminished *Haster* transcripts from the *Haster*^Stop^ allele (93% reduction, Wilcoxon rank-sum *P* = 0.02). However, we still detected abundant *Hnf1a* exon 1 transcripts from the stop allele (**Fig. 5d**). Thus, whereas deletion of the *Haster* promoter DNA caused islet-cell *Hnf1a* silencing in *cis*, this was not recapitulated by blocking *Haster* transcription. This indicates that the *Haster* promoter, but not *Haster* elongation or RNAs, is an essential positive *cis*acting element of *Hnf1a* in pancreatic islets.

### *HASTER* inhibits *HNF1A* in *cis*

Next, we examined how *HASTER* exerts negative regulation of *HNF1A*. To assess if this function also occurs in *cis*, we again examined *Hnf1a*^+/−^;*Haster*^+/−^ mice, but this time focused on liver. This showed increased HNF1A in hepatocytes, plausibly because the *Haster* deletion led to increased expression of the *Hnf1a*^+^ allele which was located on the same chromosome, and this was not compensated by *Haster* transcripts acting in *trans* from the *Hnf1a*^−^;*Haster*^+^ allele (**Fig. 5e**).Interestingly, pancreatic acinar cells showed a similar behaviour as hepatocytes in compound heterozygotes, with increased HNF1A expression (**Fig. 5b**). We also examined *Haster^+/−^ mice* bred on a hybrid strain background, and found that liver *Hnf1a* mRNA was selectively increased in *Haster* mutant alleles (**Fig. 5f**). Both findings showed that *Haster*-dependent inhibition of *HNF1A*, like its activating function, occurs in *cis*.

### *HASTER* promoter, but not its RNA, is essential for inhibition of *HNF1A*

We next examined the role of *HASTER* transcriptional elongation, RNA molecules, or its promoter in this *cis*-inhibitory function. Hybrid strain mice heterozygous for *Haster*^Stop^ showed that transcriptional blockage did not cause increased liver *Hnf1a* exon 1 transcripts in chromosomes carrying the *Haster*^Stop^ allele (**Supplementary Fig. 12**). To further examine the role of *HASTER* transcripts vs. promoters, we generated clonal EndoC-βH3 cell lines with homozygous deletions of the two *HASTER* promoters (*HASTER*^ΔP/ΔPΔ^) or a 320 bp deletion of the P1 promoter (*HASTER*^ΔP1/ΔP1Δ^) (**Supplementary Fig. 13a,b**). Both deletions caused increased *HNF1A* mRNA (**Supplementary Fig. 13a,b**), and therefore recapitulated the phenotype of mice in which *Haster* was excised after the formation of β cells (**Supplementary Fig. 8g**). To study the role of *HASTER* transcription, we targeted deactivated Cas9 to the *HASTER* transcriptional start site (CRISPRi *roadblock*^41^), or to a control intronic region located between *HASTER* and *HNF1A* promoters (**Fig. 5g**). Expectedly, this suppressed *HASTER* RNA, however this did not influence *HNF1A* mRNA or *HNF4A*, an HNF1A-dependent transcript^32^ (**Fig. 5g**). Similarly, degradation of *HASTER* nuclear transcripts using GapmeRs did not affect *HNF1A* or *HNF4A* mRNAs (**Supplementary Fig. 13c**). Conversely, CRISPR-dCas9-SAM activation of *HASTER* transcription in mouse or human β cell lines led to >5-fold higher *HASTER* RNA without changing *HNF1A* or *HNF4A* mRNAs (**Fig. 5h, Supplementary Fig. 13d**). Thus, modulation of *HASTER* transcripts or transcriptional elongation did not recapitulate the inhibitory effects of *HASTER* on *HNF1A*.

### HASTER inhibition of *HNF1A* requires HNF1A binding to *HASTER* promoter

The observation that *HASTER* transcriptional elongation was not essential was unexpected, because our genetic findings did show a tight correlation between HNF1A-dependent activation of *HASTER* transcription and negative feedback regulation of *HNF1A*. To reconcile these findings, we activated *HASTER* through lentiviral doxycycline-inducible overexpression of HNF1A (**Fig. 6a**). As in CRISPR-dCas9-SAM experiments, this led to increased *HASTER*, but this time we observed a 10-fold reduction of endogenous *HNF1A* mRNA (**Fig. 6a**). Importantly, the inhibitory effects of HNF1A overexpression where almost completely suppressed after deletion of the *HASTER* promoter region (**Fig. 6b**). Remarkably, overexpression of HNF1B, which binds to the same recognition sequences as HNF1A, and also elicited ~75% increased *HASTER* transcription, did not suppress *HNF1A* (**Fig. 6c**). These studies, therefore, demonstrated that inhibition of *HNF1A* was triggered selectively by HNF1A interactions with the *HASTER* promoter DNA, but not by various other maneuvers that influenced *HASTER* transcription.

**Fig. 6.**
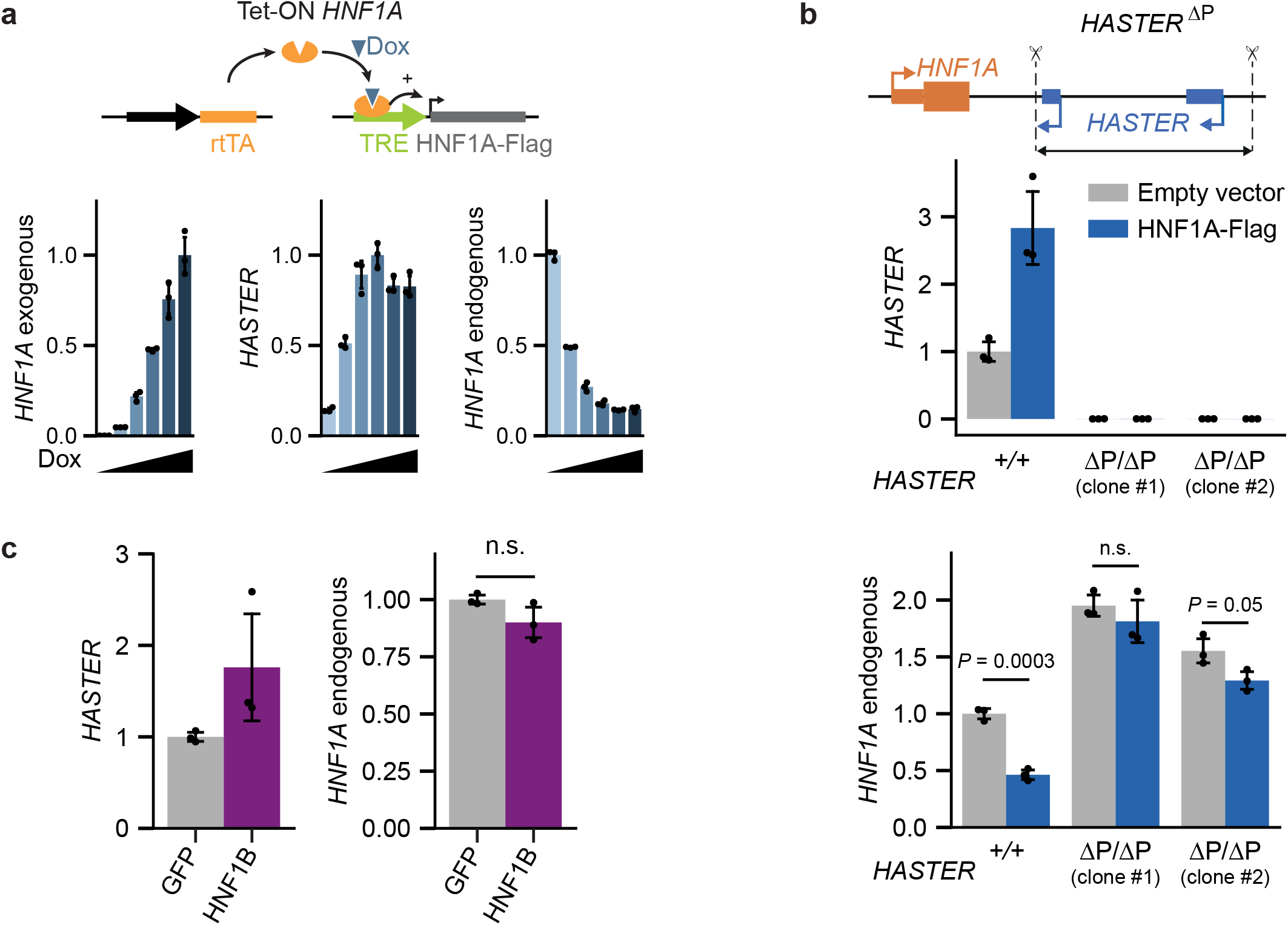
HNF1A binding to *HASTER* mediates negative regulation of *HNF1A*. **a**, Doxycycline-induced HNF1A overexpression in EndoC-βH3 cells activated *HASTER* and blocked endogenous *HNF1A* (n = 3 independent experiments). **b**, HNF1A overexpression in clonal EndoC-βH3 cell lines with homozygous deletions of both *HASTER* promoters. *HASTER* mutant clones #1 and #2 were generated using 2 independent pairs of sgRNAs; n = 3 independent experiments. **c**, *HASTER* and endogenous *HNF1A* mRNA in EndoC-βH3 nucleofected with a HNF1B or GFP expression vector (n = 3 independent experiments). **a-c**, *TBP*-normalized mean ± s.d.; two-tailed Student’s t-test.

### The *HASTER* promoter restrains *HNF1A*-enhancer spatial interactions

To further investigate how the HASTER promoter could exert negative regulation of HNF1A, we first analyzed local histone modifications. In control mouse liver, we observed, as expected, localized H3K4me3 enrichment in *Hnf1a* and *Haster* promoters. In *Haster*^LKO^, by contrast, H3K4me3 showed spreading from *Hnf1a* promoter to an intronic E enhancer (**Fig. 7a**), and H3K4me3 was significantly increased in both E and an upstream CTCF-bound C regions (t-test, P < 0.05)(**Fig. 7b**). This spreading of H3K4me3 in *Haster*^LKO^ suggested that *Haster* provides spatial insulation between *Hnf1a* promoter and the intronic E enhancer, while the small increase in H3K4me3 at E and C regions in *Haster*^LKO^ also suggested that the *Haster* deletion might cause increase proximity of E and C regions with the H3K4me3-rich *Hnf1a* promoter. We therefore hypothesized that the *HASTER* promoter could inhibit *HNF1A* by modulating 3D chromatin contacts of *HNF1A* with local regulatory elements.

**Fig. 7.**
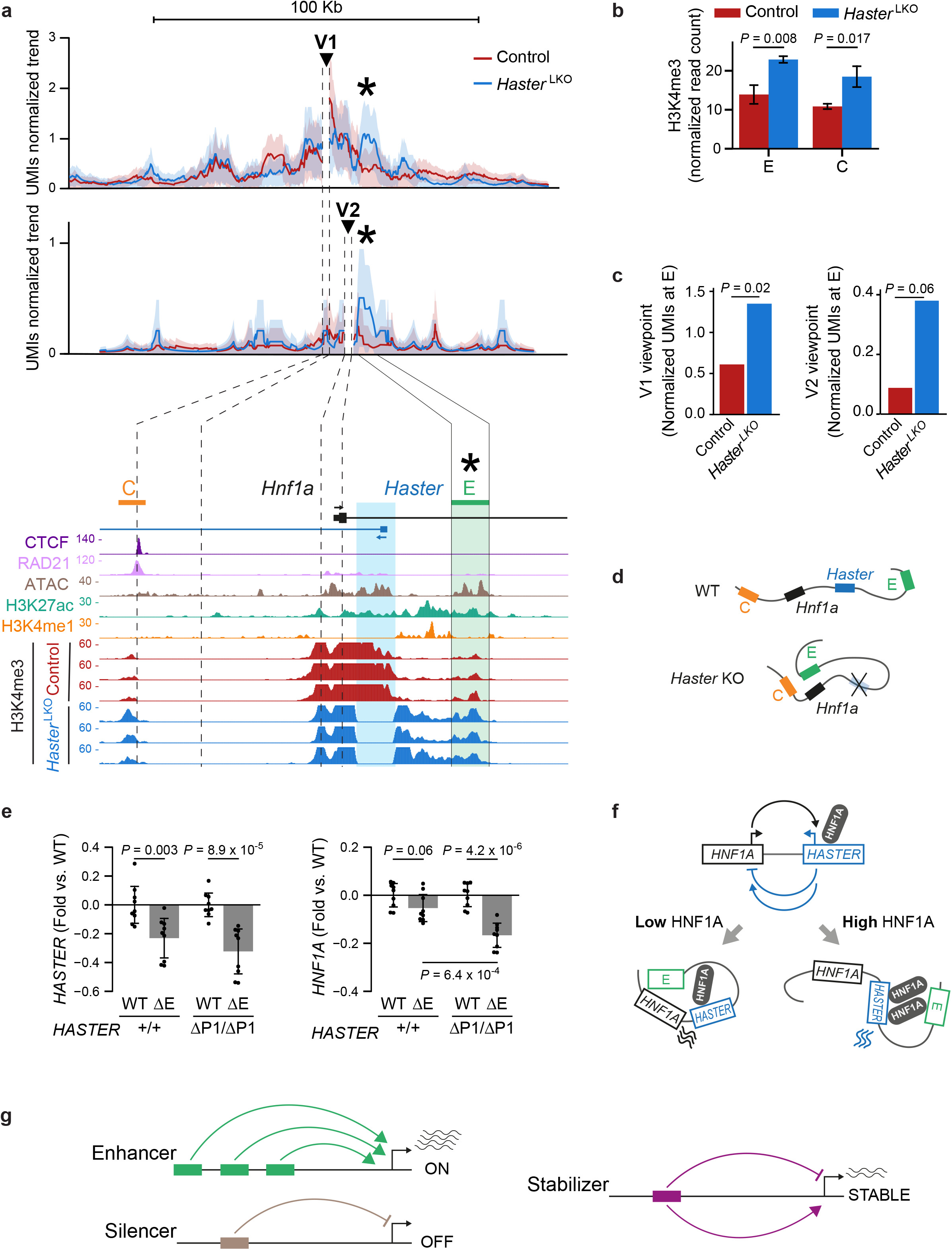
*HASTER* remodels enhancer-*HNF1A* contacts. **a**, *Haster* ^LKO^ liver shows increased contacts between *Hnf1a* upstream viewpoints (V1, V2) and the intronic E enhancer. **Top panel a**, UMI-4C profile trends for V1 (n = 6 wild type and mutant livers) and V2 (n = 3 wild type and mutant livers) viewpoints. Triangles denote viewpoints (DpnII fragment +/-1 Kb) and asterisks mark the E region. The contact profile trend reports normalized UMIs per restriction fragment in overlapping DpnII windows defined so that each include at least 5 observed molecules. Shades represent estimated binomial standard deviation. **Bottom panel a** shows H3K4me3 in liver chromatin from individual mice in the C-E region. Blue shade, region deleted in *Haster* ^LKO^. **b**, *Haster* ^LKO^ have increased H3K4me3 in C and E; n = 3 biological replicates, mean ± s.d., two-tailed Student’s t-test. **c**, UMI normalized counts at E showing increased contacts with the *Hnf1a* upstream regions (V1, V2) in *Haster* ^LKO^ liver. **d**, Schematic depicting increased interactions between *Hnf1a* and E in *Haster* ^LKO^ liver. **e**, Deletion of the E enhancer prevents increased *HNF1A* in *HASTER* deleted cells, but has limited effects on *HNF1A* in cells with intact *HASTER*. *HASTER*^+/+^ or *HASTER*^ΔP1/ΔP1^ clone #2 cells were used to create polyclonal homozygous/heterozygous E deletions (ΔE) or control WT cells lacking E deletions (sgGFP). *HASTER* and *HNF1A* RNAs are shown as fold relative to parental *HASTER*^+/+^ or *HASTER*^ΔP1/ΔP1^ cells. ΔE had significant effects on *HASTER* but not on *HNF1A* in *HASTER*^+/+^ cells, yet significantly reduced *HNF1A* mRNA in *HASTER*^ΔP1/ΔP1^ cells. Identical results were observed with E mutations in *HASTER* mutant clone #1 cells, whereas C mutations had no effects (**Supplementary Fig. 15b**). Pool of n=3 independent experiments with 3 pairs of sgRNAs for each deletion. *TBP*-normalized mean ± s.d.; two-tailed Student’s t-test. **f**, Summary model showing that *HASTER* exerts HNF1A-dependent negative regulation of *HNF1A* across tissues, and positive regulation in β cells. Left, at low HNF1A concentrations, there are unhindered interactions between the *HNF1A* promoter and E (or other local enhancers, see **Supplementary Fig. 15a**), and *HNF1A* is transcribed. Prenatally, *HASTER* itself acts as an essential enhancer in pancreatic progenitors and endocrine cells. Right, at high HNF1A concentrations, HNF1A binding to *HASTER* limits contacts between E and *HNF1A*, thereby decreasing *HNF1A* transcription. **g**, *HASTER* is a *cis*-acting stabilizer that prevents target overexpression and silencing. This function is distinct from enhancers or silencers, which drive cell- and stage-specific transcription or silencing, respectively.

To test this, we performed quantitative chromatin conformation capture (UMI-4C)^42^ experiments. Mouse *Hnf1a* and *Haster* promoters as well as intronic E enhancer are all located within ~7 Kb, a linear distance that limits the analysis with chromatin conformation methods. We thus selected two viewpoints to study 3D chromatin interactions at the *Hnf1a* locus: one viewpoint ~6 Kb upstream of *Hnf1a*, near the CTCF-bound C site (viewpoint 1), and another that contained the *Hnf1a* promoter (viewpoint 2)(**Fig. 7a**). UMI-4C experiments from *Haster*^LKO^ vs. control liver (n = 6 per genotype) showed that the *Haster* deletion caused >2-fold increased contacts between both *Hnf1a* upstream regions and the intronic E enhancer (V1, Chi-square test for pooled UMI-4C libraries, *P* = 0.02)(**Fig. 7a,c, Supplementary Fig. 14**). Thus, the analysis of two viewpoints showed consistent changes in interactions between the *Hnf1a* upstream region and the intronic E enhancer in *Haster*^LKO^ (**Fig. 7d**). These observations therefore suggested that *HASTER* limits 3D contacts between *HNF1A* and the intronic E enhancer, thereby limiting *HNF1A* transcription. Consistent with this model, deletion of the intronic E enhancer prevented the increase in *HNF1A* mRNA that results from *HASTER* deletions, but had insignificant effects in cells with intact *HASTER* (**Fig. 7e, Supplementary Fig. 15a,b**). Taken together, these experimental models show that *HASTER*-dependent negative feedback of *HNF1A* occurs through a *cis* function of the *HASTER* promoter that does not require *HASTER* transcription. It is instead mediated by HNF1A binding to the *HASTER* promoter, which insulates *HNF1A* from cis-acting positive regulatory elements (**Fig. 7f)**.

## Discussion

These studies uncover a *cis*-regulatory element that provides both activating and inhibitory functions to ensure that *HNF1A* is expressed at appropriate concentrations in each cell type (**Fig. 7f**). They also show that HNF1A binding to a 320 bp lncRNA promoter sequenc, but not the transcriptional elongation or RNA products are essential for this function. The dual nature of this regulatory element is most compellingly illustrated by the coexistence of HNF1A silencing or overexpression in β cells from *Haster*-deficient mice. This stabilizing function is fundamentally different from *cis*-acting transcriptional enhancers or silencers that provide spatiotemporal ON or OFF switches (**Fig. 7g**).

Our findings show that *HASTER*-dependent feedback is a critical determinant of how HNF1A selects genomic high-affinity recognition sites in different cell types. Interestingly, a few lncRNAs have been recently shown to exert negative modulation, through different mechanisms, on the heart transcription factor gene *Hand2*^12,43^, the c-MYC oncogene^12^, or *CHD2*^15^. All such genes, *HAND2*, *MYC*, *CHD2*, as well as *HNF1A*, share in common that they are haploinsufficient genes encoding for transcriptional regulators^15,18–44,45^, while c-MYC, HAND2, and HNF1A have been used in misexpression systems for lineage reprogramming – a feature of transcription factors that can act on repressed chromatin^36,46^. These examples, and most clearly *HASTER*’s dual function, suggest that the principal function of a group of lncRNA units may be to stabilize dosage sensitive transcriptional regulators that have a capacity to transform cell-specific chromatin landscapes, and thus require controlled intracellular concentrations to ensure the selectivity of their genomic targets.

*HASTER* also provides an example of a genetic defect in a mammalian lncRNA that causes diabetes mellitus. Remarkably, the main manifestation of germline *Haster* mutations was β cell dysfunction and diabetes, which was due to its pancreatic islet-specific role to maintain *Hnf1a* transcription. Interestingly, *HNF1A* heterozygous mutations also primarily cause selective β cell dysfunction and only subclinical alterations in other cell types^18^, whereas homozygous *Hnf1a* mutations cause severe liver and renal dysfunction, growth retardation, diabetes, and embryonic lethality^21,24^. The discovery of a lncRNA stabilizer of *HNF1A*, therefore, provides insights to dissect cell-specific genetic mechanisms underlying *HNF1A* haploinsufficient diabetes, and to modulate *HNF1A* function in β cells.

Finally, this finding has general implications for our understanding of noncoding genome defects in disease. Transcriptional enhancers often form clusters that provide robustness to genetic disruption^47,48^, whereas our findings indicate that the 320 bp *HASTER* promoter region lacks functional redundancy. This warrants a need for careful examination of lncRNA elements in genomes of disease cohorts.

## Acknowledgements

This research was supported by grants from Medical Research Council (MR/L02036X/1), Wellcome Trust (WT101033), European Research Council Advanced Grant (789055), Spanish Ministry of Science and Innovation (BFU2014-54284-R, RTI2018-095666-B-I00), and National Institute for Health Research (NIHR) Imperial Biomedical Research Centre. Work in CRG was supported by the CERCA Programme, Generalitat de Catalunya and Centro de Excelencia Severo Ochoa (SEV-2015-0510) and support from the Spanish Ministry of Science and Innovation to the EMBL partnership. We thank the University of Barcelona School of Medicine animal facility, Center of Genomic Regulation and Imperial College London Genomics Units, and the Imperial College High Performance Computing Service. We thank Klaus Kaestner (University of Pennsylvania), Maureen Gannon (Vanderbilt University), and David Tuveson (Cold Spring Harbor Labs) for Cre lines. We thank Juan Valcarcel and Thomas Graf (Center for Genomic Regulation) for comments on the manuscript, Vanessa Grau and Charlotte Roth for expert technical support, and members of the Ferrer lab for valuable discussions.

## Author Contributions

A.B., J.F. conceived, coordinated and supervised the study. A.B. performed cell based and computational studies, supervised mouse analysis. A.B., M.A.M., J.G. performed analysis of mouse mutants. M.G.D.V. and P.R. designed and created cell models, R.G.F. performed FISH with supervision from A.B. A.B. and I. M-E performed UMI-4C studies. A.B., Aina Bernal and D.B. performed stem cell studies. A.B. designed mouse models with input from J.F., S.O., P.V. S.O. and P.V. created homologous recombinant mice. H.H. participated in single cell genomics. A.B., J.F. wrote the manuscript with input from the remaining authors.

## Notes

### Competing Interest Statement

The authors have declared no competing interest.

