## Supplementary material for "*HASTER* is a transcriptional stabilizer of *HNF1A*": Methods

### Methods, Data and code availability

#### Animal studies

Animal experimentation was carried out in compliance with the EU Directive 86/609/EEC and Recommendation 2007/526/EC regarding the protection of animals used for experimental and other scientific purposes, enacted under Spanish law 1201/2005, and all experiments were approved by the Institutional Animal Care Committees of the University of Barcelona and Parc de Recerca Biomedica de Barcelona. *Haster f* (LoxP) and *stop* (transcriptional termination) alleles were generated in C57Bl/6N JM8.F6 embryonic stem cells (ES cells) by homologous recombination. Briefly, mouse ES cells were electroporated with a linearized targeting plasmid containing the *Haster* promoter flanked by LoxP sites, or a transcription termination (3 x SV40 polyA) signal downstream of the *Haster* promoter, as well as a PGK/Neomycin selection cassette flanked by FRT recombination sites (**Supplementary Fig. 4**). Constructs were linearized by PmeI and SacII for the conditional allele, and by PacI and PmeI for the stop allele. Electroporated mouse ES cells were selected for the targeting cassette with Geneticin. Clones were analyzed by Southern blot, and correctly targeted clones were injected into C57BL/6BrdCrHsd-Tyrc morulae (E2.5) to create chimeric mice that transmitted the recombined allele through the germ line. The PGK/Neo cassette was excised by crossing heterozygous mice with Tg(CAG-Flp) mice. Mice were bred on C57Bl/6 backgrounds unless otherwise specified.

To excise *Haster* in pancreatic epithelial cells $Haster^{+/f}$ mice were crossed with *Pdx1*-Cre mice (Tg(Pdx1-Cre)6Tuv)[1]. Constitutive excision in $\beta$ cells was achieved with $Ins1^{Cre}$ mice, in which Cre is inserted in the *Ins1* locus ($Ins1^{tm1.1(cre)Thor}$)[2]. Inducible excision in $\beta$ cells was achieved with the *Pdx1*-$CreER^{TM}$ transgene (Tg(Pdx1-Cre/Esr1*)1Mga)[3] after two oral deliveries of 40 µg of Tamoxifen (Merk), spaced by 4 days, in 10- to 13-week-old mice, and analyzed 12 weeks later. Deletion at the outset of liver development was achieved with *Alb*-Cre mice, in which Cre is driven by an albumin promoter and alphafetoprotein enhancer (Alb Tg(Alb1-cre)1Khk)[4]. *Haster* germ-line deletions were generated by breeding $Haster^{+/f}$ mice with Tg(EIIa-cre) mice[5]. $Hnf1a^{+/-}$ mice have been described[6]. Genotype was confirmed by PCR using the primers provided in **Supplementary Table 4**.

Lines with LoxP alleles without Cre, Cre lines without LoxP alleles and wild-type littermates served as controls, as indicated for each experiment. Experimental cohorts of young adult mice were maintained on a 12-hour light/dark cycle with free access to water and standard mouse chow and housed in specific pathogen-free barrier facilities. Prior to decapitation, mice were anesthetized using isoflurane (Zoetis).

#### Glucose tolerance

Animals were fasted overnight and received intraperitoneal glucose injections (2 g/Kg). Blood was collected at indicated times for measuring glucose using GlucoMen aero2k meter (Menarini Diagnostics).

For meal-tests, male age-matched animals were fasted for 16 hr overnight and blood glucose and insulin were measured before and 1 hr after refeeding.

Plasma insulin was quantified using the Ultra Sensitive Mouse Insulin ELISA kit (Crystal Chem) following manufacturer's recommendations and were read using Infinite M Plex (Tecan) plate reader. Standard curves were fitted using quadratic polynomial regression. Each assay was performed in duplicate using 5 µL of plasma from mouse tail. Insulin plasmatic concentration was determined by taking the mean of duplicates.

**Islet isolation**
Islet isolation was performed as previously described[7]. Briefly, the pancreas was inflated with an ice-cold collagenase P solution (1 mg/mL in Hanks' balanced salt solution (HBSS) buffer, Roche) injected through the main pancreatic duct. The pancreas was then dissected and incubated at 37ºC for 8 min with continuous agitation. The pancreatic tissue was disaggregated by gentle suction through a needle, and cell suspension was washed 4 times with cold HBSS-0.5% BSA. The islet pellet was resuspended in 7 mL Histopaque containing a 7:3 ratio of pre-cooled Histopaque 1077 (Merck) and Histopaque 1119 (Merck), and 7 mL of HBSS-0.5% BSA was layered on top of the Histopaque. The gradient was centrifuged at 950 g for 20 mins at RT. The interphase containing the islets was collected, washed 3 times with HBSS-0.5% BSA and islets were finally further enriched by gentle aspiration with a pipette under a stereomicroscope. Islets were allowed to recover for 2 days in culture in RPMI containing 11 mM glucose supplemented with 10% fetal calf serum and penicillin-streptomycin (1:100; Invitrogen) at 37°C and 5% $CO_2$.

**Vector construction**
To achieve high Cas9 and dCas9 expression in EndoC-βH3 cells, we generated lentiviral vectors with the human insulin promoter driving expression of Cas9 or dCas9, in addition to a U6-driven sgRNA (pLV-hIP-Cas9-BSD and pLV-hIP-dCas9-BSD). EF1a core promoter and puromycin resistance of lentiCRISPRv2 vector (plasmid #52961, Addgene) were replaced by the human insulin promoter and blasticidin-S deaminase (BSD) using Gibson Assembly (NEBuilder® HiFi DNA Assembly Master Mix, NEB, E2621X) to generate the pLV-hIP-Cas9-BSD. The human insulin promoter (343bp) was amplified from EndoC-βH3 genomic DNA and the BSD from the lenti-dCAS-VP64-Blast (plasmid # 61425, Addgene). Cas9 from the pLV-hIP-Cas9-BSD was replaced by dCas9 using Gibson Assembly to generate the pLV-hIP-dCas9-BSD, and dCas9 was amplified from the pSp-dCas9-2A-GFP[8].

sgRNAs for CRISPRi *road block* were designed within 100 bp downstream of the transcriptional start site defined by CAGE in human islets. 20 nt sgRNAs were designed using Cas-Designer (http://www.rgenome.net/cas-designer/) and cloned into expression vectors as described[9]. Briefly, oligonucleotides (Thermo Fisher) containing sgRNA sequences flanked by compatible overhangs with the destination vectors were phosphorylated with T7 polynucleotide kinase (NEB) and annealed. Oligonucleotide duplexes were ligated into BbsI- or BsmBI-digested destination vectors. Ligated constructs were transformed into Stbl3 chemically competent *E. coli* and clones were checked by Sanger sequencing. For deletions, sgRNA pairs were cloned as described[10]. Briefly, a fragment containing the scaffold of the sgRNA1 and the H1 promoter of the sgRNA2 were amplified from the pScaffold-H1 donor (plasmid #118152, Addgene) with primers containing the protospacer of the sgRNA1 and 2 as well as a BbsI restriction site. The PCR fragment was digested with BbsI and ligated into the destination vector.

A TetOn HNF1A lentiviral vector (pLenti-CMVtight-HNF1A-FLAG-Hygro) was built by cloning the human HNF1A-FLAG fragment into the pLenti CMVtight Hygro DEST (plasmid # 26433, Addgene). The rtTA was expressed from a lentiviral vector (pLV-rtTA-zeo) that was built by amplifying the UbC promoter and rtTA-Advance cassette from the pHAGE-TRE-dCas9-KRAB (#50917, Addgene), and cloned into a lentiviral backbone, upstream to a 2A-ZeocinR cassette. Sequences of sgRNAs and vector used in this study are listed in **Supplementary Table 5**.

**Cell culture**

EndoC-βH3 cells[11] were maintained on 2 µg/mL fibronectin- and 1 % ECM-coated plate in Dulbecco's Modified Eagle Medium (DMEM) low glucose (1 g/L) supplemented with sodium pyruvate (Thermo Fisher), 2% albumin from bovine serum fraction V (Roche), 1% heat-inactivated FBS (Labtech), 2 mM L-glutamine, 5.5 µg/mL human transferrin, 1 mM sodium pyruvate 10 mM nicotinamide, 6.7 ng/ml sodium-selenite, 50 µM β-mercaptoethanol, 100 U/mL penicillin and 100 µg/mL streptomycin. DMEM was substituted by Advance DMEM-F12 (Thermo Fisher) and FBS was omitted for the TetOn-HNF1A EndoC-βH3 cell line, and during the expansion of EndoC-βH3 clones.

293FT cells (Thermo Fisher) were maintained in DMEM, 10% heat-inactivated FBS, 0.1 mM MEM non-essential amino acids, 2 mM L-glutamine, 1 mM sodium pyruvate, 500 µg/ml geneticin, 100 U/mL penicillin and 100 µg/mL streptomycin.

MIN6 cells[12] were maintained in DMEM 4.5 g/L glucose, 15% heat-inactivated FBS, 50 µM β-mercaptoethanol and gentamicin 50 µg/mL.

**Cell-based knockdowns, overexpression and CRISPR-based modifications**

LNA GapmeRs (Exiqon, **Supplementary Table 6**), pEZY3-HNF1A-Flag, pcDNA3.1-HNF1B and CRISPR/Cas9 plasmid vectors were nucleofected in EndoC-βH3 cells using a Nucleofector B2 (Lonza), with Amaxa cell line nucleofection kit V (Lonza) and program G-017. We used 2 million cells and 10 µg plasmid DNA per nucleofection for CRISPR/Cas9 deletion or 1 million cells with 250 pg of LNA GapmeRs. Cells were harvested 72 hr after GapmeR or plasmid nucleofection.

CRISPRi and CRISPR-SAM lentiviral particles were produced as previously described[10]. 293FT cells were seeded at 75,000 cells/cm$^2$ in T75 flasks, and 24 hr later transfected with CRISPRi or CRISPRa vectors and third generation packaging plasmids pMDLg/pRRE, pRSV-Rev and pMD2.G (#12251, #12253 and #12259, Addgene) with PEI-Pro (Polyplus-transfection) following manufacturer's instructions, in antibiotic-free media using a 1:1 ratio of total DNA µg to µl of PEI-Pro. Medium was replaced by 9 ml fresh 293FT antibiotic-free media 18 hours post-transfection and lentiviral particles were collected 72 hr post transfection. Immediately after collection, supernatants were centrifuged for 5 min at 400g and filtered using Steriflip-HV, 0.45µm, PVDF filters (Millipore). Supernatant was supplemented with 1 mM $MgCl_2$ and treated with 1 µg/mL DNaseI (Roche) for 20 minutes at 37ºC. Virus particles were then concentrated with 1:3 v/v of Lenti-X Concentrator (Clontech) at 4ºC overnight. On the following day, virus particles were collected by cold centrifugation for 45 min at 1500g, resuspended in 100 µL of PBS, aliquoted and stored at -80ºC until use. Transduction was carried out with 10 µL virus for 400,000 cells in 1 mL. Antibiotic selection was started 3 days after transduction with 8 µg/mL blasticidin, 100 µg/mL hygromycin or 200 µg/mL zeocin for EndoC-βH3 cells.

A CRISPR-SAM cell line was established by successive transduction of lentivirus dCAS-VP64_Blast (#61425, Addgene) and MS2-P65-HSF1_Hygro (#61426, Addgene). For activation of specific genes, the cells were transduced with lentivirus sgRNA(MS2)_zeo (#61427, Addgene) expressing a sgRNA targeting the desired gene promoter. CRISPR/Cas9 and CRISPRi road block experiments were performed by transduction of lentivirus hIP-Cas9-BSD (pLV-hIP-Cas9-BSD plasmid) and hIP-dCas9-BSD (pLV-hIP-dCas9-BSD plasmid) respectively.

For CRISPR/Cas9 clonal deletions, EndoC-βH3 cells were nucleofected with pSpCas9(BB)-T2A-HygR (#118153, Addgene) containing the sgRNA. 24 hr after nucleofection, the cells were selected using Hygromycin B (200 µg/mL, ThermoFisher, 10687010) for 3 days. After two weeks, the cells were seeded at low density (2-9 cells/$cm^2$) in Advance DMEM-F12 based EndoC-βH3 medium. EndoC-βH3 clones were hand-picked and transferred into 96-well plates. After genotyping, selected clones were expanded and maintained in DMEM-based EndoC-βH3 medium with 1% FBS.

A doxycycline-inducible HNF1A EndoC-βH3 cell line was established by successive transduction of EndoC-βH3 cells with a rtTA-2A-ZeoR-expressing lentivirus (pLV-rtTA-zeo) and TRE-HNF1A-FLAG lentivirus carrying the hygromycin resistance gene (pLenti-CMVtight-HNF1A-FLAG-Hygro). Cells were exposed to doxycycline (0, 25, 50, 100, 200 and 400 ng/mL) for 24 hr before analysis. Endogenous HNF1A mRNA was detected with oligonucleotides that recognize a 3' untranslated region of HNF1A that is not present in HNF1A-FLAG, whereas exogenous HNF1A-FLAG was detected with oligonucleotide primers that recognize FLAG (**Supplementary Table 4**).

**hESC genome editing**

H9 hESCs were maintained in mTeSR1 medium (StemCell Technologies, 85870) on Matrigel (Corning, 356231) coated plate. For nucleofection, cells were dissociated with Accutase (Merk) for 8 min at 37ºC, then diluted in 10 µM Y27632 (Merk) mTeSR1, centrifuged at 110g for 3 min and resuspended in 10 µM Y27632 mTeSR1. $10^6$ cells were nucleofected using the Human Stem Cell Nucleofector® Kit 2 (Lonza; program G-017) with 5 µg of pSpCas9(BB)-2A-puro (#62988, Addgene) expressing 2 sgRNAs. After nucleofection, the cells were transferred to a 12-well containing 1 mL of 10µM Y27632 mTeSR1. After 24h, the cells were selected for puromycin resistance by replacing the medium with fresh 10 µM Y27632 mTeSR1 containing 0.5 µg/mL puromycin for 24h. After selection, the hESCs were cultured in mTeSR1 medium without Y27632. After 2 passages, cells were dissociated and plated at low density. Isolated clones were transferred and maintained in 96-well plates until genotyping for homozygous deletions.

**hESC differentiation**

H9 clones carrying the different deletions were differentiated to hepatocytes using a protocol adapted from Hannan *et al.*[13]. Cells were seeded at 300,000 cells per 24-well in 10 µM Y27632 mTeSR1 and differentiation was started after 24 hr. The following mediums were used for differentiation: (a) S1 medium[14] was prepared with MCDB 131 medium (ThermoFisher, 10372019) supplemented with 8 mM D-(+)-Glucose (Merk, G7528), 2.46 g/L NaHCO3 (Merk, S3817), 2% Bovine Serum Albumin Fraction V (Roche, 10735078001), 1:50.000 Insulin-Transferrin-Selenium-Ethanolamine (ITS -X) (ThermoFisher, 51500056), 2 mM GlutaMAX (ThermoFisher, 35050061) and 0.25 mM L-Ascorbic acid (Merc; A4544); (b) RPMI/B27 medium: RPMI 1640 Medium, GlutaMAX™ Supplement (ThermoFisher, 61870010) supplemented with B-27™ Supplement (ThermoFisher, 17504044) and MEM Non-Essential Amino Acids Solution (ThermoFisher, 11140035); (c) Hepatocyte growth medium (HGM): HBM Basal Medium (Lonza, CC-3199) supplemented with 3.75 g/mL Bovine Serum Albumin Fraction V, 250 µg/mL L-Ascorbic acid, 10 µg/mL Holo-transferin (Merk, T0665), 0.5 µg/mL Hydrocortisone (Merk, H0888), 5 µg/mL Human Insulin and 10 ng/mL EGF (R&D Systems, 236-EG-200). During differentiation, the medium was changed every day or every 2 days after day 11, using the following mediums: Day 1, S1 medium with 100 ng/ml ActivinA (R&D Systems, 338-AC-050) and 3 μM Chir99021 (Tocris, 04-0004); Day 2 and 3, S1 medium with 100 ng/ml ActivinA; Day 4 to 6, RPMI/B27 medium with 50 ng/ml ActivinA; Day 7 to 10, RPMI/B27 medium with 20 ng/mL BMP-4 (R&D Systems, 314-BP-010) and 10 ng/mL FGF10 (Source BioScience, ABE1324); Day 11 to 25, HGM medium with 30 ng/mL Oncostatin M (OSM) (R&D Systems, 295-OM-010) and 50 ng/mL HGF (Peprotech, 100-39). Definitive endoderm stage was reached at day 4, anterior endoderm stage at day 7, hepatoblast stage at day 11 and hepatocyte stage at day 26.

Differentiations to pancreatic progenitor and endocrine cells were performed using a modification of a published protocol[15]. Dissociated hESC were seeded at 2 million cells per 35 mm well coated with Matrigel in 5 μM Y27632 E8 medium (ThermoFisher, A1517001). Differentiation was started the following day after washing the cells once with 1XPBS: Definitive endoderm induction: day 0, MCDB131 + 2mM GlutaMax (ThermoFisher, 35050038) + 1.5 g/L NaHCO3 + 0.5% BSA fraction V (Lampire, 7500804) + 10 mM final glucose + 100 ng/ml Activin A (Qkine, QK001) + 3 μM CHIR99021 (Tocris, 4423); day 1, same as day 0, reducing CHIR99021 to 0.3 uM; day 2, same as day 1 with no CHIR99021; Stage 2, Posterior Foregut induction, day 3-5, MCDB131 + 2 mM GlutaMax + 1.5 g/L NaHCO3 + 0.5% BSA fraction V + 10 mM final glucose + 0.25 mM ascorbic acid (Sigma, A4544) + 50 ng/mL FGF7 (Genscript, Z03407-1); Stage 3, Pancreatic Endoderm induction, day 6-7, MCDB131 + 2 mM GlutaMax+ 2.5 g/L NaHCO3 + 2% BSA fraction V + 10 mM final glucose + 0.25 mM ascorbic acid + 50 ng/mL FGF7 + 0.25 μM SANT1 (Sigma, S4572), 1 μM Retinoic acid (Sigma, R2625), 100 nM LDN193189 (Selleckchem, S2618), 1:200 ITS-X (ThermoFisher, 51500-056), 200 nM TPB (SantaCruz, sc-204424); Stage 4, Pancreatic Progenitor induction, day 8-11, MCDB131 + 2 mM GlutaMax + 2.5 g/L NaHCO3 + 2% BSA fraction V + 10 mM final glucose + 1:200 ITS-X +0.25 mM ascorbic acid + 2 ng/mL FGF7 + 0.25 uM SANT1+ 0.1 uM Retinoic acid + 200 nM LDN + 100 nM TPB + 100 ng/mL EGF (Peprotech, AF-100-15) + 10 mM Nicotinamide (Sigma, N0636) + 10 ng/mL Activin A + 10 uM Y27632. Cells were dissociated with TrypLE and seeded in AggreWell™400 plates (Stem Cell Technologies, 34425) on day 10; Stage 5, Endocrine Progenitor induction, day 12-15, MCDB131 + 2 mM GlutaMax+ 1.5 g/L NaHCO3 + 2% BSA fraction V + 20 mM final glucose + 1:200 ITS-X + 10 μg/mL Heparin (Sigma, H3149) + 10 μM Zinc Sulfate (Sigma, Z0251) + 0.05 μM Retinoic acid + 0.25 μM SANT1 + 100 nM LDN + 1 μM T3 (Sigma, T6397) + 100 nM GSiXX (Merck, 565789) + 10 μM ALK5inhII (Selleckchem, S7223) + 20 ng/mL Betacellulin (Peprotech, 100-50); Stage 6, Endocrine Cell induction, day 16-23, cell aggregates were transferred to ultra low attachment 6-well plates and placed in an orbital shaker at 100 rpm, MCDB131 + 2 mM GlutaMax+ 1.5 g/L NaHCO3 + 2% BSA fraction V + 20 mM final glucose + 1:200 ITS-X + 10 μg/mL Heparin + 10 μM Zinc Sulfate + 100 nM LDN + 10 μM ALK5inhII + 1 μM T3 + 100 nM GSiXX.

**RT-qPCR**

RNA was prepared using RNeasy mini kit (Qiagen) including DNAse I (Qiagen) treatment. Total RNA was retrotranscribed using SuperScript III (ThermoFisher) and random hexamers (ThermoFisher) following manufacturer's protocol. Quantitative PCR was performed with Universal Probe Library assays (UPL, Roche). Reactions were carried out in duplicates, in a QuantStudio 12K Flex (Applied Biosystems) with 1x TaqMan Fast Advanced Master Mix (Thermo Fisher), 1 µM forward and reverse primers and 250 nM of UPL probe, or 1x TaqMan assay. Quantification was performed using the standard curve method and the mean quantity of the duplicates was used to calculate the gene expression. Gene expression was normalized by a reference gene, *TBP* or *RPLP0*. Primers and ULP probes are listed in **Supplementary Table 4**.

**Single molecule Fluorescence In Situ Hybridization (smFISH)**
smFISH was performed as described[16]. A set of 48 probes (**Supplementary Table 7**), coupled with Quasar570 (548/566) or Quasar670 (647/670), was designed for each transcript (Stellaris RNA FISH probes, Biosearch Technologies). EndoC-βH3 cells were grown on fibronectin/ECM-coated (2 µg/mL fibronectin, 1% ECM, Merk) coverslips. Cells were fixed in 4% formaldehyde for 2 min, washed with 1X PBS and permeabilized with 70% ethanol at 4ºC for at least an hour. Probe hybridization was performed overnight at 37°C in the dark with hybridization buffer (10% formamide and 100 mg/ml of dextran sulfate in 2X SCC) containing 12.5 µM of probes. The following day, cells were washed for 30 minutes at 37°C with a buffer containing 10% formamide in 2X SCC, followed by another 30 minutes incubation with a solution of 5 ng/ml of 4',6-diamidino-2-phenylindole (DAPI). Finally, coverslips were mounted using Vectashield Hard set mounting media. Acquisitions were performed at the Facility for Imaging by Light Microscopy, Imperial College London on inverted widefield microscope (Zeiss Axio Observer Inverted Widefield Microscope) with LED illumination. Z-stacks acquisitions were taken with the 63x objective every 0.5 µm from a 40 µm total depth, deconvoluted (Huygens software) and the maximal projections of the whole stacks were used for the counting process. Between 8 to 12 fields were recorded and counted for each sample.

**Immunofluorescence**
Embryos and adult tissues were collected at indicated times and processed for immunofluorescence analysis of paraffin embedded pancreas as previously described[17]. Briefly, tissues were fixed in 4% paraformaldehyde overnight at 4ºC, then washed in PBS before paraffin embedding. 4 µm sections were deparaffinized in xylene and rehydrated with ethanol series. Sections were incubated for 30 min at room temperature in antibody diluent (DAKO Corporation) with 3% normal serum from the same species as the secondary antibody and incubated overnight at 4 °C with primary antibody, then overnight at 4 °C with secondary antibody, and finally DAPI stained and mounted with Dako Fluorescence Mounting Medium (Molecular Probes, S3023). The following primary antibodies were used: monoclonal rabbit anti-HNF1A (1:400, D7Z2Q, Cell Signaling Technology), guinea pig anti-insulin (1:200, A0564, Dako), quinea pig anti-glucagon (1/1000, 4030-01F, Millipore), rat anti-Cytokeratin 19 (1/100, Hybridoma Bank, TROMA III-c), goat anti-PDX1 (1:200, AF2419, R&D Systems), mouse anti-Nkx6.1 (1:200, F55A10, Hybridoma Bank). Images were acquired using Leica TSE confocal microscope for immunofluorescence performed on tissue, and using Leica DMi8 for immunofluorescence performed on cell lines.

**Protein isolation and western blotting**
Proteins were extracted from frozen mouse livers with 9M Urea solution. Quantification was performed using Microplate BCA protein assay kit (ThermoFisher,

23250) according to the manufacturer's protocol. Western blot was performed with 20 µg of total proteins on a 4-12% Bis Tris gel (ThermoFisher, NP0335BOX**)**. Proteins were detected using β-tubulin (Cell signalling, 2146) and HNF1A (Cell Signaling, 89670) antibodies.

**Cellular fractionation**
Cellular fractionation was performed as described[18]. 5 million EndoC-βH3 cells were incubated 5 min on ice in 200 µL cold Lysis Buffer (10 mM Tris-HCl pH 7.5, 0.05% IGEPAL, 150 mM NaCl, 100 U/mL SuperaseIn (ThermoFisher)). The cell lysate was layered over 2.5 volumes of chilled Sucrose Solution (10 mM Tris-HCl pH 7.5, 0.05% IGEPAL, 150 mM NaCl, 24% sucrose, 100 U/mL SuperaseIn) and centrifuged 10 min at 15,000g, 4ºC. The supernatant was kept as the "cytoplasmic fraction". The pellet was washed with 500 µL Wash Buffer (1 mM EDTA in PBS pH 7.5) and centrifuged 10 min 15,000g, 4ºC. Then, the pellet was resuspended in 100 µL cold Glycerol Buffer (20 mM Tris-HCl pH 7.5, 75 mM NaCl, 0.5 mM EDTA, 0.85 mM DTT, 1x protease inhibitor cocktail (Roche), 50% glycerol, 100 U/mL SuperaseIn), and 100 µL cold Nuclei Lysis Buffer (10 mM HEPES pH 7.5, 1 mM DTT, 7.5 mM $MgCl_2$, 0.2 mM EDTA pH 8, 0.3 mM NaCl, 1 M urea, 1% IGEPAL) was added to the nuclei suspension, vortexed and left on ice for 2 min. Nuclear lysate was centrifuged for 2 min at 15,000g, 4ºC and supernatant were collected as nucleoplasmic fraction. The pellet ("chromatin fraction") were washed with 500 µL Wash Buffer, and resuspended in 300 µL Chromatin DNase Buffer (20 mM Tris-HCl pH 7.5, 50 mM KCl, 4 mM $MgCl_2$, 0.5 mM $CaCl_2$, 2 mM TCEP (Merk), 1x protease inhibitor cocktail, 0.4% sodium deoxycholate, 1% IGEPAL, 0.1% N-lauroylsarcosine). 15 µL of murine RNase inhibitor (NEB) and 30 µL TURBO DNase (Ambion) was added and the reaction was incubated 20 min at 37ºC. DNase was inactivated by adding 12.5 µL of 25x Stop Solution (250 mM EDTA, 125 mM EGTA). Proteins were digested with 7.5 µL of Proteinase K (Ambion) for 1h at 37ºC. RNA from the different fractions was purified using Zymo RNA Clean & Concentrator-25 kit (Zymo Research).

**3' RACE**
3' RACE was performed as described[19]. 240 ng of human islet RNA were retrotranscribed with $Q_T$ primers (**Supplementary Table 4**) using Superscript III. Nested PCRs were performed with Q5 polymerase (NEB). For the first PCR, we used 1:20$^{th}$ of cDNA with a gene-specific forward primer 1 and a $Q_O$ reverse primer, and for the second PCR, 1 µL of a 1:5 dilution of the 1$^{st}$ PCR with a gene-specific forward primer 2 and a $Q_I$ reverse primer (primers in **Supplementary Table 4**). The resulting fragments were gel-purified (QIAquick, Qiagen), cloned and sequenced.

**ChIP**
Liver tissue was collected after perfusion of ice-cold PBS. Tissue was then minced with a razor blade in ice-cold PBS. Minced liver tissue (100 mg) or 100-500 mouse islets were incubated with 1% formaldehyde (Agar Scientific) for 10 min at RT. The reaction was stopped by addition of 1:10$^{th}$ of 1.25 M glycine for 5 min at RT, pelleted at 4°C for 3 min at 800g and washed twice with PBS. Aliquots containing 20 mg of initial liver tissue or all processed islets were snap-frozen and stored at -80°C until use. Crosslinked samples were lysed using ice-cold Lysis Buffer (2% Triton X-100, 1% SDS, 100 mM NaCl, 10 mM Tris-HCl pH 8, 1 mM EDTA pH 8, 1x protease inhibitor cocktail) for 15-20 minutes on ice. Chromatin was sonicated using a S220 Focused-ultrasonicator (Covaris), with the following settings: 2% Duty Factor, 105W

Peak Incident Power, 200 Cycles per Bust for a duration of 16 minutes. Sheared chromatin was centrifuged at full speed 10 min 4°C to remove debris and insoluble chromatin, and the supernatant was transferred to a fresh low-binding tube. For liver, chromatin equivalent to 5 µg of DNA was used for one histone mark ChIP and 10 µg for transcription factor ChIP. Chromatin was diluted 4 times with ChIP Dilution Buffer (0.75% Triton X-100, 0.1% Na-deoxycholate, 140 mM NaCl, 50 mM HEPES pH8, 1 mM EDTA, 1x protease inhibitor cocktail) and 5 % was used as input. 30 µL of Dynabeads Protein G (ThermoFisher) were blocked with BSA overnight at 4ºC. 10 µL of HNF1A antibody (Cell Signaling Technology, D7Z2Q), 2 µg of H3K27ac antibody (Abcam, ab4729) and 2 µg of H3K4me3 antibody (Merk, 15-10C-E4), or 2 µg of H3K4me1 antibody (Abcam, ab8895) were added to the samples and incubated overnight rotating at 4°C. 30 µl of magnetic beads were added to the samples and rotated at 4°C for 2h.

For ChIP-qPCR, antibody-incubated samples were washed with Low Salt Wash Buffer (1% Triton X-100, 0.1% SDS, 150 mM NaCl, 20 mM Tris-HCl pH 8, 2 mM EDTA pH 8), High Salt Wash Buffer (1% Triton X-100, 0.1 % SDS, 500 mM NaCl, 20 mM Tris-HCl pH 8, 2 mM EDTA pH 8), LiCl Wash Buffer (0.25 M LiCl, 1% IGEPAL, 1% sodium deoxycholate, 10 mM Tris-HCl pH 8, 1 mM EDTA pH 8) and 3 times with TE. Elution was performed with 200 µL of Elution Buffer (1% SDS, 0.1M $NaHCO_3$) for 30 min at RT. Samples were placed on a magnet and the supernatant was transfer to a new tube. 1 µL RNase A (ThermoFisher) was added to the eluate and incubated for 30 min at 37ºC. Reverse crosslink was performed by adding 8 µL of 5M NaCl and 3 µL of proteinase K (ThermoFisher) and incubated 1 hr at 55˚C, 1200 rpm, then overnight at 65˚C, 1200 rpm. DNA was purified using MinElute PCR Purification Kit (Qiagen). Quantitative PCR reactions were carried out in duplicates as described for RT-qPCR. Allele-specific qPCR were performed using Custom TaqMan SNP Genotyping Assays. Enrichment was subsequently normalized by the input. Primers, ULP probes and TaqMan assays are listed in **Supplementary Table 4**.

For ChIPmentation, washes and tagmentation were performed as previously reported[20]. Antibody-incubated samples were washed twice with RIPA-LS (10 mM Tris-HCl pH 8, 140 mM NaCl, 1 mM EDTA pH 8, 0.1% SDS, 0.1% sodium deoxycholate, 1% Triton X-100), twice with RIPA-HS (10 mM Tris-HCl pH 8, 500 mM NaCl, 1 mM EDTA pH 8, 0.1% SDS, 0.1% sodium deoxycholate, 1% Triton X-100), twice with RIPA-LiCl (10 mM Tris-HCl pH 8, 250 mM LiCl, 1 mM EDTA pH 8, 0.5% IGEPAL, 0.5% sodium deoxycholate) and once 10 mM Tris-HCl pH 8. Beads were resuspended in 20 µL of tagmentation solution (10 mM Tris-HCl pH 8, 5 mM $MgCl_2$, 10% v/v dimethylformamide) containing 1 µL Tn5 (Illumina) and incubated at 37ºC for 10 min. The reaction was stopped by adding 1 mL ice-cold RIPA-LS buffer and incubated 5 min on ice. Beads were then washed twice with RIPA-LS, twice with TE, and resuspended in Elution buffer (10 mM Tris-HCl pH 8, 5 mM EDTA pH 8, 300 mM NaCl, 0.4% SDS). Proteinase K was added to the elution and incubated 1h at 55˚C, 1200 rpm, then overnight at 65˚C, 1200 rpm. DNA was purified using MinElute PCR Purification Kit (Qiagen). To estimate the number of cycles required for the amplification of the libraries, 2 µL of the elution was used for Syber Green qPCR, using Kapa HiFi polymerase (Kapa Biosystems). Libraries were amplified from 20 µL of elution with Kapa HiFi polymerase and Nextera custom primers (**Supplementary Table 4**), for Ct value roundup + 1 cycles. DNA clean-up was performed with 1.8x volume and size selection with 0.65x volume of AMPureXP beads (Beckman Coulter). Libraries were sequenced on HiSeq 2500 using 1 x 50 bp reads.

**ChIP-seq analysis**
ChIP-seq reads were aligned with Bowtie2 (v.2.3.5) on GCRm38 genome. Alignment statistics are listed in **Supplementary Table 8**. Multi-mapped reads were discarded. Reads mapping to the ENCODE blacklisted regions were removed using BedTools (v.2.27.1). Duplicated reads were removed using Picard (v.2.6.0). Peak calling was performed using MACS2 (v.2.1.1) with a minimum FDR cut-off (-q) of 0.05. The --broad flag was used for histone marks peak calling. MACS2 bdgcmp function was used to generate the local Poisson test $-\log_{10}$ *P*-values. P-value BedGraphs were converted to bigWig using bedGraphToBigWig UCSC tool. Differential binding analysis was performed using the Bioconductor R package DiffBind (v.2.8.0) on all peaks called in at least 2 samples from any genotypes and using normalized read coverage from triplicates. Significant binding differences were determined at FDR $q \leq 0.05$. HNF1A neo-binding sites were defined as peaks observed in at least 2 *Haster* KO samples ($q \leq 0.05$), without significant peaks called by MACS2 in any control sample, as well as an average $\log_2$ normalized ChIP read counts $\leq 2$ in control samples. Activated promoters were similarly defined as H3K4me3 enrichment peaks detected in at least 2 *Haster* KO ($q \leq 0.05$) but not in any control sample, with $\log_2$ normalized ChIP read counts < 2 in controls, significant differential H3K4me3 enrichment ($q \leq 0.05$) and positive fold change in *Haster* vs. controls. Coverage was calculated using deepTools (v.3.0.2) computeMatrix and the average of the 3 replicates was calculated for each bin. Peak intersections were performed with pybedtools (v.0.8.0). Mouse liver ChIP-seq for CTCF was from GSE29184[21]; RAD21 from GSE102997[22]; FOXA2, CEBPB and HNF4A from GSE57559[23]; PPARA from GSE108689[24]; RXR from GSE35262[25]; GATA4 from GSE49132[26].

**Motif analysis**
Motif analysis of known and *de novo* transcription factor binding site motifs was performed using Homer (v.3.12). All analyses were performed on the full length of the merge between overlapping consensus peaks defined by DiffBind, merging peaks from any sample that showed minimum 1 bp overlap. Enrichment analysis of *de novo* transcription factor motifs was performed with the findMotifsGenome.pl command on consensus peaks defined by DiffBind, for a length of 8, 10 and 12 bp on masked mm10 genome.

**ATAC-seq**
We used previously reported ATAC-seq reads from mouse liver and kidney[27]. Reads were trimmed to remove Illuman Nextera adaptors using Trim Galore and aligned with Bowtie2 (v.2.3.5) on GCRm38 genome. Multi-mapped and duplicated reads were removed using Picard (v.2.6.0). Mitochondrial and ENCODE blacklisted region reads were discarded. For visualization, MACS2 bdgcmp function was used to generate the local Poisson test $-\log_{10}$(*P*-value) bedGraphs. BedGraphs were converted to bigWig using bedGraphToBigWig UCSC tool. Coverage was calculated using deepTools (v.3.0.2) computeMatrix for 1 kb windows with 10 bp bins.

**RNA-seq**
RNA from islets or liver was quantified with Qubit (ThermoFisher) and the quality was verified with Bioanalyzer (Agilent). Libraries were prepared with TruSeq Stranded mRNA Library kit and sequenced on Hiseq4000 using 2 x 75 bp reads. Reads were aligned to the GCRm38 genome with STAR (v.2.3.0) aligner for

coverage visualization. Transcript level quantification was performed with Salmon (v.0.11) using GENCODE GCRm38 vM18 annotation (**Supplementary Table 8**). Gene level normalization and differential expression was performed using the Bioconductor R package DESeq2 (v.1.24.0), using adjusted $P \leq 0.05$ as a cut off for differentially expressed genes. Fold changes were adjusted using lfcShrink using the apeglm option[28].

Differential expression of transcripts associated with HNF1A binding sites was performed as follow. *De novo* transcripts from *Haster*$^{LKO}$ and control liver were assembled from RNA-seq using StringTie (v.2.0). Transcripts from *Haster*$^{LKO}$ and control replicates were merged in a single GTF file using gffcompare (v.0.10.1). Transcript quantification and differential transcript expression were performed using Salmon and DESeq2 as described above, using the merged *Haster*$^{LKO}$ and control liver transcriptome as reference. Transcripts with low abundance (mean normalized TPM < 3) were discarded. To define transcripts with an HNF1A-bound promoter, a minimum of 1 bp overlap between the TSS and an HNF1A peak was required.

Human (SRX218942)[29], chicken (SRX2704301) and X. tropicalis (SRX2704321)[30] liver RNA-seq reads were aligned on GCRh38 (hg19), galGal5 and XenTro9 genome respectively. Mouse kidney (SRX2370375) and small intestine (SRX2370402) RNA-seq reads were obtained from the Mouse ENCODE project[31], and were aligned to the GCRm38 (mm10) genome.

RNA-seq of hPSC differentiated to β cells were obtained from GSE140500[32].

**Allele-specific RNA-seq**

Stranded total RNA libraries from C57BL/6;PWK/PhJ F1 liver and islets were sequenced on HiSeq 2500 using 2 x 125 bp reads. Reads were aligned by Star (v.2.7.6) using the WASP algorithm[33], and reads that aligned to a different genomic region after swapping the C57BL/6 variant to the PWK/PhJ variant were discarded. PWK/PhJ SNPs were obtained from the Mouse Genome Project (version 5)[34]. ASEReadCounter (GATK v.4.1.9.0) was used to count the reads overlapping *Hnf1a* and *Haster* PWK/PhJ exonic SNPs.

**HNF1A-regulated gene set**

To define a high-confidence gene set that is positively regulated by HNF1A in islet cells we intersected two sets: (a) 115 genes that showed downregulation in Affymetrix array analysis of islets from *Hnf1a*$^{-/-}$ islets[35], and (b) 570 genes that showed increased expression (adjusted $P \leq 0.05$ and mean normalized expression > 500) after CRISPR-SAM activation of *Hnf1a* in MIN6 mouse β cells. Briefly, the MIN6-SAM cell line was generated by successive transduction of MIN6 mouse β cells with lentivirus dCAS-VP64_Blast (#61425, Addgene) followed by blasticidin selection (1 µg/mL) and lentivirus MS2-P65-HSF1_Hygro (#61426, Addgene) followed by hygromycin selection (100 µg/mL). MIN6-SAM cells were subsequently transduced with the sgRNA expressing vector (lenti sgRNA-(MS2)-zeo (#61427 , Addgene). RNA from technical triplicates of 2 independent *Hnf1a* activating sgRNAs and 2 independent control sgRNAs was extracted. RNA-seq and differential expression analysis were performed as described above. In total, 21 genes showed concordant downregulation and upregulation in the two models, respectively.

**Gene Set Enrichment Analysis (GSEA) and Enrichr**

Enrichment of predefined sets of genes among genes that change in experimental conditions was performed using GSEAPreranked (v.6.0, GenePattern)[36] on genes ranked by KO vs. control fold change, using default parameters over 10000 permutations to calculate false discovery rates. Enrichments of functional gene annotations in differentially expressed genes were performed with Enrichr[37].

**Tissue-specificity Z-score**

A gene tissue-specificity z-score in each *Hnf1a*-expressing tissue was calculated for each gene by taking the average normalized gene expression in the tissue minus the mean of all *Hnf1a*-expressing tissues, divided by the standard deviation of all *Hnf1a*-expressing tissues, as described[38].

**Single-cell RNA-seq**

Cultured mouse islets were dissociated with Accutase (Merk) for 15 min at 37ºC. Islet cell suspensions were centrifuged at 600g 3 min and resuspended in cell culture medium with DAPI before FACS sorting to remove dead cells and doublets. After sorting, cells were centrifuged at 600g for 3 min and resuspended in PBS/0.04% BSA. Single-cell libraries were generated with 10x Genomics Chromium Single Cell 3′ Reagent Kit v3 following the manufacturer's instructions. Libraries were sequenced on HiSeq 4000.

**Single cell RNA-seq analysis**

Read alignments and UMI count were performed using CellRanger (v.3.0.2) using mm10 reference genome. Subsequent analyses were carried out with Seurat (v.3.0.1). Cells with <500 genes or >5% mitochondrial genes were filtered out (**Supplementary Table 8**). UMI counts were normalized using SCtransform function[39]. To define shared population between Controls and KOs, we performed an integrated analysis on the 3 control and 3 KO datasets [40]. Briefly, the 3000 most variable genes were used to find anchors (SelectIntegrationFeatures function using 50 dimensions). The first 50 principal components were used for t-SNE projection (RunTSNE function) and clusters were defined by graph-based unsupervised clustering (FindClusters function) with a resolution of 0.5.

Differential expression was performed with the FindMarkers function (min.pct = 0.1) for all combination of controls and KOs. Wilcoxon Rank Sum *P*-values from the different combinations were combined using Fisher's method. Only genes with consistent positive or negative fold change across all control/KO combinations and with a combined $P \leq 0.05$ were considered differentially expressed. All genes differentially expressed in endothelial cells were discarded from the analysis. For differential expression analysis, all β cell clusters with more than 250 cells were grouped in a single cluster named β-cluster.

Seurat objects were exported as loom using the as.loom function of the loomR (v.0.2.0.1) library. Data visualization was performed with python (v.3.7.3) and loompy (v.2.0.16), numpy (v.1.15.4), pandas (v.0.25.0), matplotlib (v.3.1.0) and seaborn (v.0.9.0) libraries. Statistics were computed using scipy (v.1.1.0).

**UMI-4C**

UMI-4C was performed as described[41], with minor modifications. Liver tissue from three samples per genotype were collected and crosslinked with 2% formaldehyde for 10 min as described above for ChIP. Frozen pellets of ~$10^7$ cells were thawed on ice and resuspended in 5 mL of Cold Lysis Buffer (50 mM Tris-HCl pH 7.5, 150 mM NaCl, 5 mM EDTA, 1% Triton X-100, 0.5% IGEPAL, 1x protease inhibitor cocktail). After isolation, the nuclei were resuspended in 650 µL of nuclease-free water, 60 µL

DpnII buffer, 15 µL 10% SDS and incubated at 37ºC, 1h, 900 rpm, with an additional hour after addition of 75 µL of 20% Triton X-100. The chromatin was digested at 37ºC, 900 rpm, for 24 hr using 600U of DpnII (R0543L, NEB), and enzyme was inactivated by incubating at 65ºC for 20 min. Ligation was performed in a final volume of 7 mL with 60 U of T4 DNA ligase (Promega) and incubated at 16ºC overnight. The efficiency of the digestion and ligation was assessed by gel electrophoresis. Chromatin was reverse crosslinked with 30 µL Proteinase K (10 mg/mL), overnight at 65ºC, followed by 45 min of incubation with 30 µL RNaseA (10 mg/mL) at 37ºC. The DNA was purified by phenol-chloroform extraction followed by ethanol precipitation, and resuspended in 10 mM Tris-HCl pH 8. 10 µg of DNA was sonicated using a S220 Focused-ultrasonicator (Covaris), to obtain 400-600 bp fragments. DNA was end-repaired with 10 µL NEBNext End Repair Mix (NEB, E6050L) in 200 µL final volume, incubated 30 min at 20ºC, purified with 2.2x AmpureXP beads (Beckman Coulter), and eluted in 10 mM Tris-HCl pH 8. A-tailing was performed with 200 U Klenow fragment (NEB, M0212M) in 100 µL 1X NEBbuffer 2 with 1nM dATP. 5' ends were dephosphorylated at 50ºC for 60 min with 20U calf intestinal alkaline phosphatase (NEB, M0290S). The DNA was then cleaned with 2x AmpureXP beads. Adapters were ligated with 0.4 µM Illumina-compatible forked indexed adapters (**Supplementary Table 4**) and 10 µM quick ligase (NEB, M2200) in 160 µL 1x quick ligase buffer (NEB, M2200) for 15 min at 25ºC. DNA was denatured at 95ºC 2 min and cleaned with 1x AmpureXP beads. To generate UMI-4C libraries, two nested PCRs were performed using Kapa HiFi polymerase (Kapa Biosystems) with 0.4 mM final concentration of primers. The 1$^{st}$ PCR used the upstream bait primer (**Supplementary Table 4**) and the Illumina universal primer 2, and amplification was performed for 20 cycles. The DNA was cleaned with 1x AmpureXP and used for the 2$^{nd}$ PCR with the downstream bait primer (**Supplementary Table 4** and the Illumina universal primer 2 for 16 cycles. After the 2$^{nd}$ PCR, the DNA was cleaned, and size selected with 0.7x AmpureXP beads. The size distribution of the libraries was controlled by Bioanalyzer and the libraries were quantified by qPCR with KAPA library quantification kit (Roche, 07960166001). Libraries from controls and *Haster*$^{LKO}$ mice were sequenced on HiSeq 2500 using 2 x 125bp reads.

**UMI-4C-Seq analysis**

UMI-4C datasets were analysed using umi4cPackage (v 0.0.0.9000) as described[41]. To increase UMIs complexity, fastq files from sequenced libraries were initially pooled by genotype. Paired-end sequencing reads were demultiplexed using fastq-multx from ea-utils (v.1.3.1). Reads were aligned and the number of UMIs extracted using *p4cCreate4CseqTrack* function. A window of 1kb around the viewpoint was removed from the analysis. 4C contact profiles from the KO and control animals were normalized for UMI coverage using *plotCompProf* function and an adaptative smoothing method that controls window size so that no less than 5 molecules are included in each window. Assessment of differential contacts between KO and WT 4C profiles in genomic regions of interest within a 0.5 Mb window surrounding the viewpoint was carried out using *p4cIntervalsMean* through a chi-square test of normalized molecule counts.

**Data availability**

Raw sequence reads from RNA-seq, scRNA-seq, and ChIP-seq are available from Arrayexpress, under accession number (pending). Processed data files are provided as Supplementary Data Sets and deposited at https://www.crg.eu/en/programmes-groups/ferrer-lab#datasets (pending).

### Code availability

All custom code used to generate the data in this study is available upon reasonable request.
