## Supplementary Figures and Legends for "*HASTER* is a transcriptional stabilizer of *HNF1A*"

### Supplementary Fig. 1

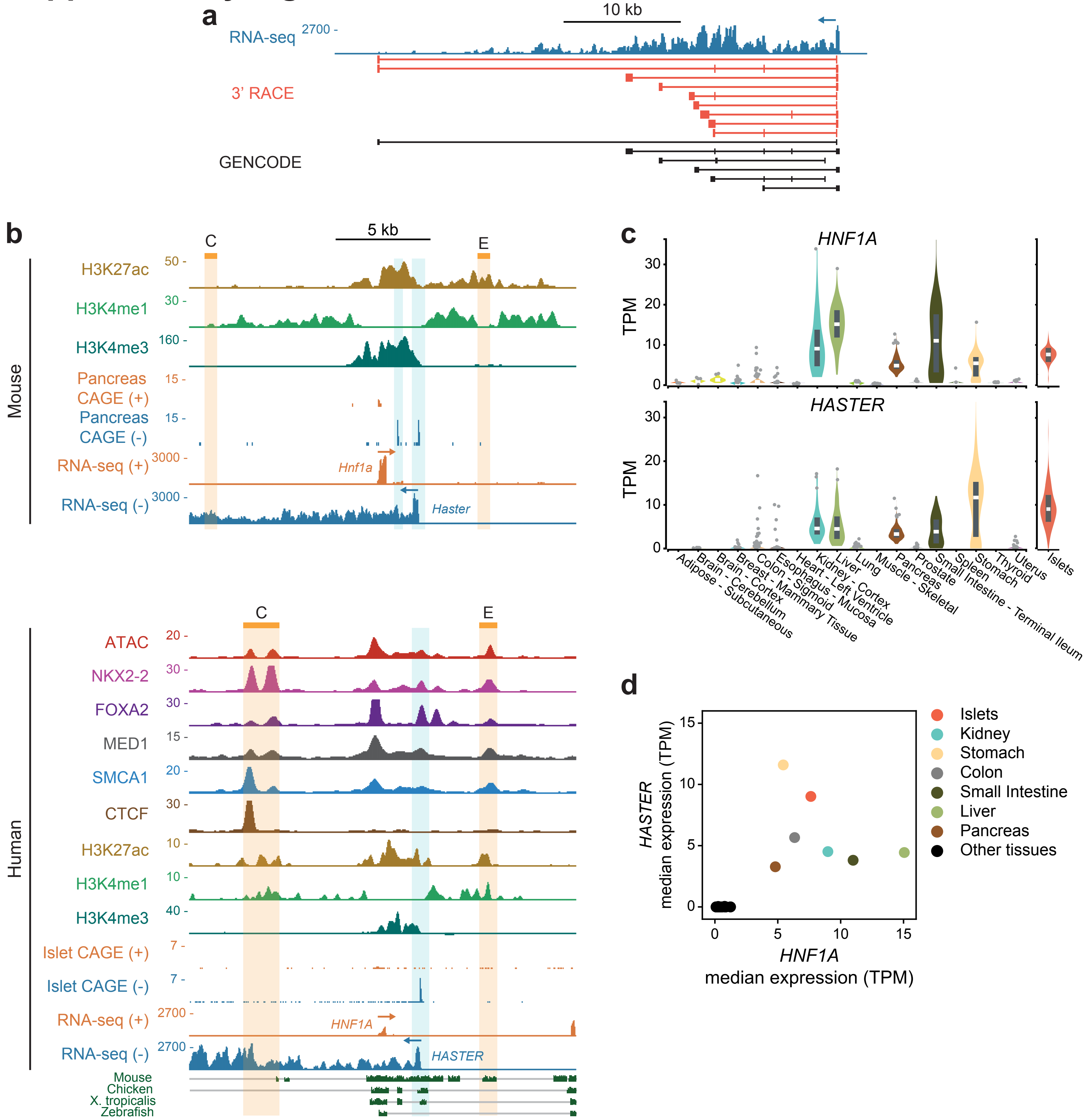


**Supplementary Fig. 1. *HASTER* gene structure and expression. a**, *HASTER* isoforms detected by 3' Rapid Amplification of cDNA Ends. **b**, Chromatin and RNA maps in mouse (top) and human islets (bottom). Human CAGE is from islets, and mouse CAGE is from pancreas. The two mouse *Haster* transcriptional start sites are highlighted in blue, although only one transcriptional origin is apparent in human islets. The E1 islet enhancer, and CTCF bound C region, both of which are bound by islet transcription factors, are highlighted in beige. **c**, *HNF1A* and *HASTER* expression across GTEx human tissues and human islets. Boxes show median and interquartile ranges. **d**, *HASTER* and *HNF1A* median transcript levels across tissues are negatively correlated, with the exception of whole pancreas.

### Supplementary Fig. 2

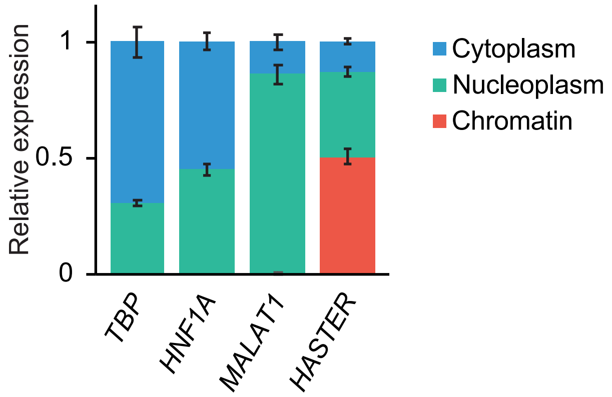


**Supplementary Fig. 2. *HASTER* localizes to the nucleus.** Relative subcellular expression of *HASTER* lncRNA in EndoC-βH3 cells, compared to control mRNAs (*TBP* and *HNF1A*) and the nuclear lncRNA *MALAT1*. Mean ± s.d., n = 3 biological replicates.

### Supplementary Fig. 3

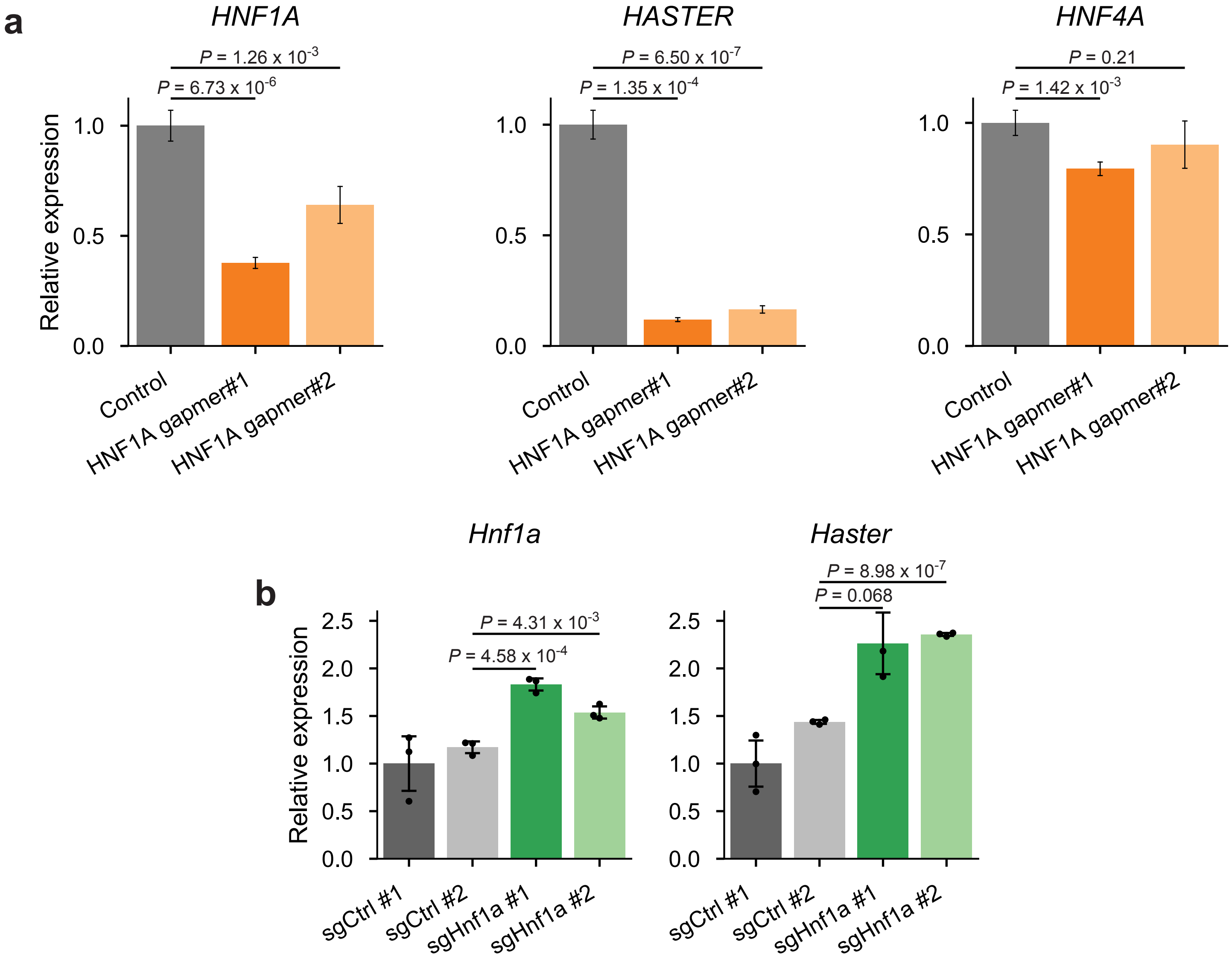


**Supplementary Fig. 3. *HASTER* is sensitive to decreased or increased *HNF1A* expression. a**, LNA GapmeR knock-down of *HNF1A* in human EndoC-βH3 β cells led to decreased *HASTER*, and minor changes in other HNF1A-dependent genes. n = 3 nucleofections, *TBP*-normalized mean ± s.d.; two-tailed Student's t-test. **b**, CRISPR-SAM activation of *Hnf1a* in mouse MIN6 β cells. n = 3 lentiviral transductions, representative of 2 independent experiments. *Tbp*-normalized mean ± s.d.; two-tailed Student's t-test.

### Supplementary Fig. 4

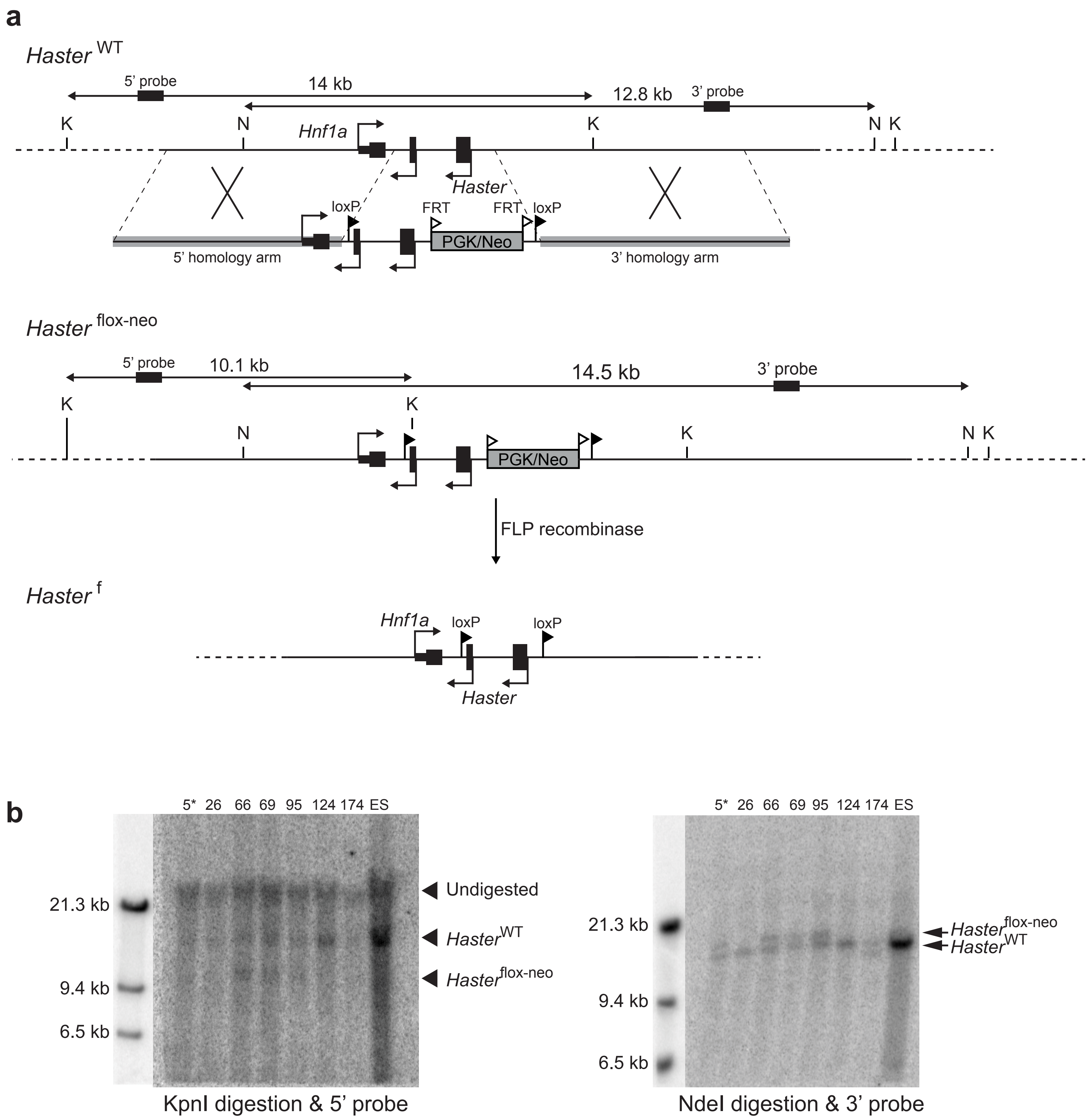


**Supplementary Fig. 4. Conditional mouse *Haster* allele. a**, Schematic of the targeted allele, digestion fragments and probes used for Southern blot analysis of different alleles. **b**, Southern blot with KpnI (left) and NdeI (right) digestion. Asterisk, Clone 5 was selected to establish the line. K, KpnI; N, NdeI; ES, parental embryonic stem cell (C57BL/6).

### Supplementary Fig. 5

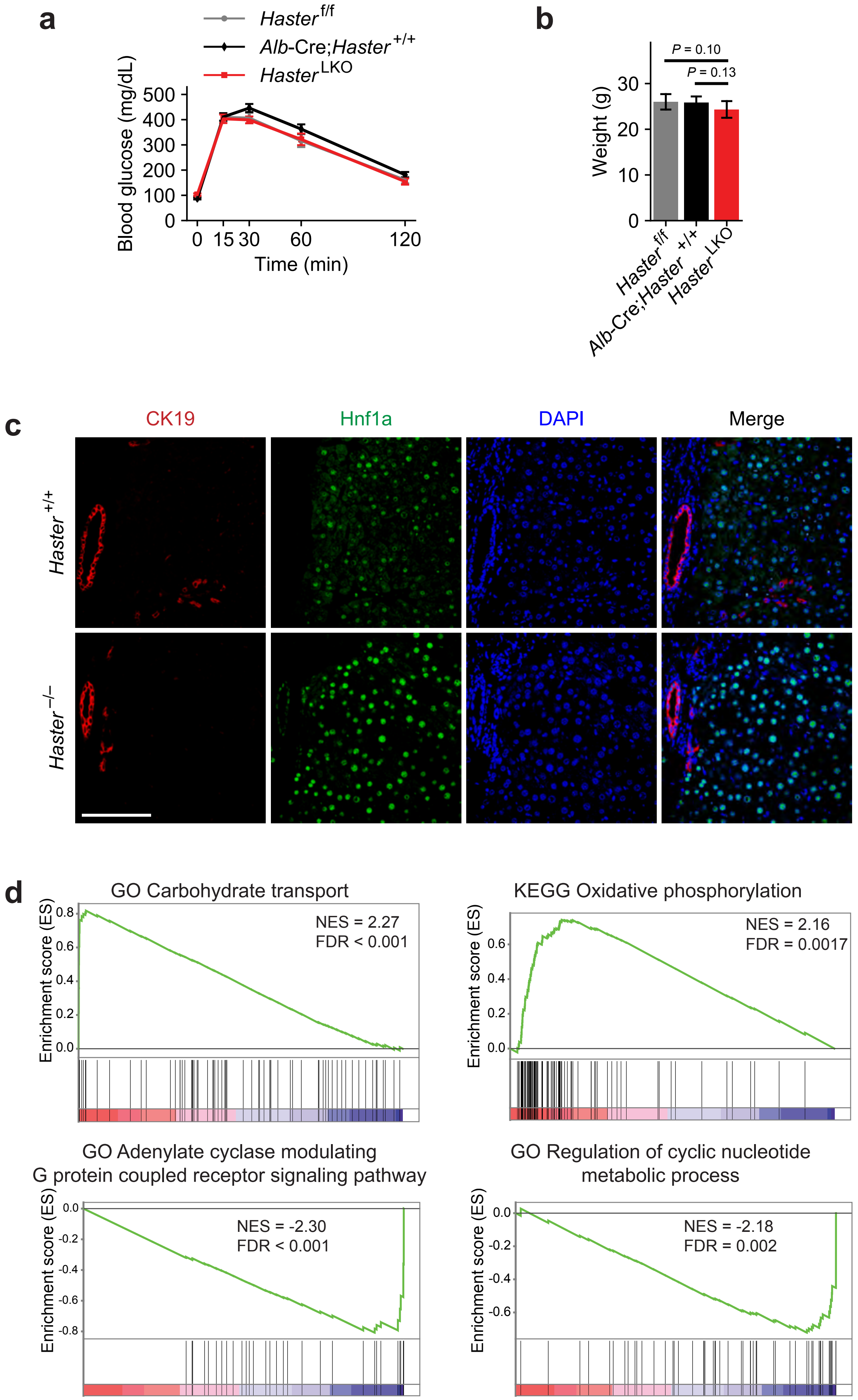


**Supplementary Fig. 5. Phenotypic analysis of *Haster* deletion in liver. a**, Intraperitoneal glucose tolerance test in *Haster*$^{LKO}$ and control 8-week-old mice. Mean ± s.e.m. **b**, Body weight at 8 weeks for *Haster*$^{LKO}$ and controls. Mean ± s.d.. **a**,**b**, n = 9 *Haster*$^{LKO}$, n = 6 *Alb*-Cre;*Haster*$^{+/+}$ and n = 7 *Haster*$^{f/f}$, two-tailed Student's t-test. **c**, Immunofluorescence showing HNF1A overexpression in *Haster*$^{-/-}$ liver. Scale bar, 100 µM. **d**, GSEA displaying the enrichment of functional annotations in *Haster*$^{LKO}$ upregulated (top panel) and downregulated (bottom panel) genes.

### Supplementary Fig. 6

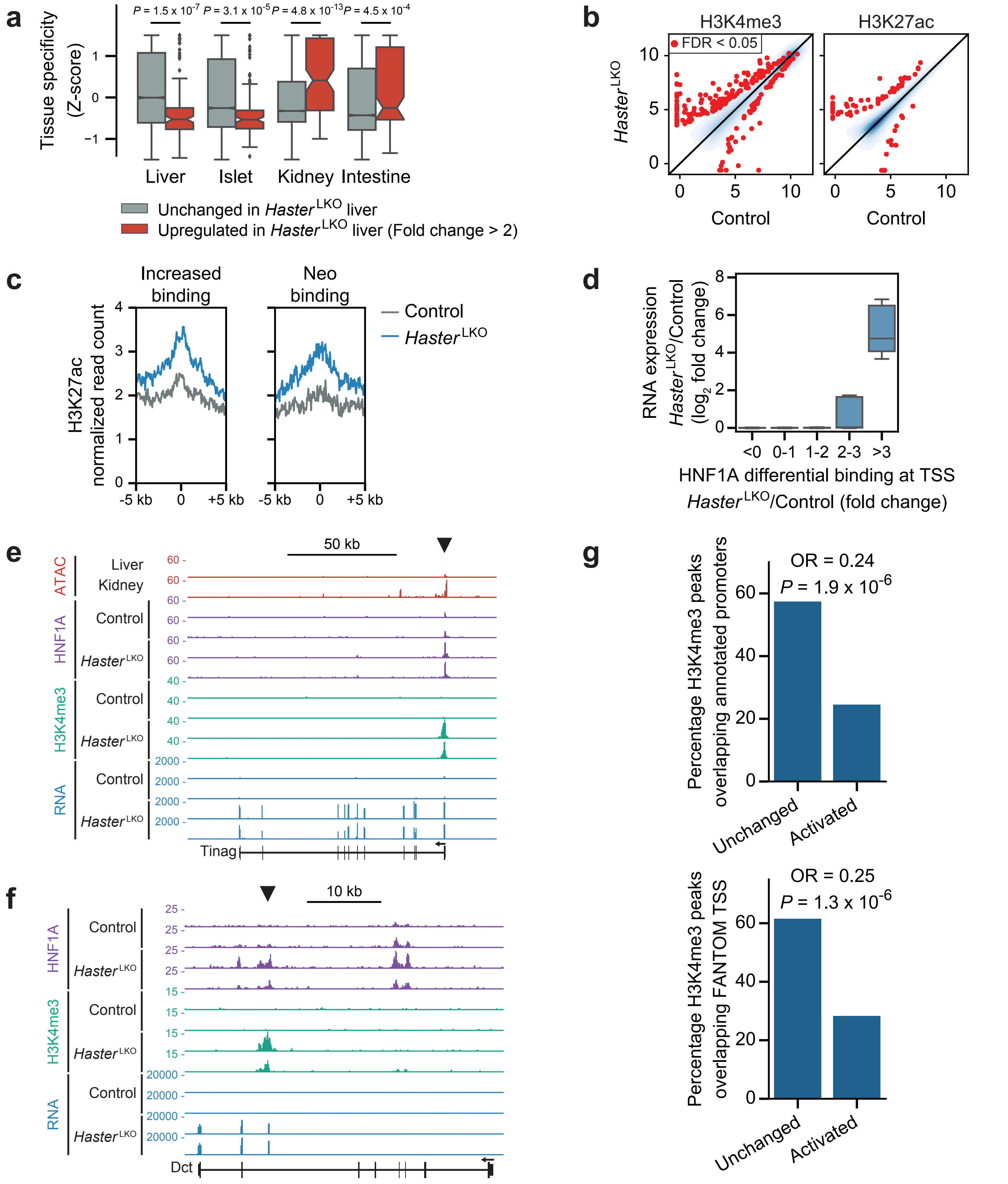


**Supplementary Fig. 6. *Hnf1a* upregulation perturbs HNF1A binding selectivity. a**, Tissue specificity of gene expression across HNF1A-expressing tissues for genes upregulated in *Haster*LKO liver. To quantify tissue specificity, for each gene and tissue we calculated a Z-score that represents the deviation of expression in that tissue relative to the average from all tissues. Box plots show medians and interquartile ranges. Wilcoxon rank-sum *P*-values. **b**, Liver H3K4me3 and H3K27ac in *Haster*LKO and control liver (log2 normalized ChIP-seq read count; n = 3 mice per genotype). Red, differential H3K4me3 or H3K27ac sites (FDR ≤ 0.05); blue, kernel density of differential H3K4me3 or H3K27ac sites with FDR > 0.05. **c**, H3K27ac at HNF1A-bound regions in *Haster*LKO and controls (average of n = 3 mice per genotype). **d**, RNA fold change in *Haster*LKO vs. control liver of HNF1A-bound promoters for the different categories of HNF1A binding in *Haster*LKO liver. Lines are median and interquartile ranges. **e**, Examples of HNF1A neo-binding sites that lead to ectopic promoter and gene activation in *Haster*LKO liver. **f**, Ectopic activation of an intragenic promoter in *Haster*LKO liver (n = 2 mice per genotype). **g**, Activated genomic regions that are bound by HNF1A and become active promoters in *Haster*LKO, but are inactive in control liver, overlap less frequently with annotated promoter and FANTOM5 CAGE transcriptional start sites, compared with HNF1A-bound active promoters in control liver (unchanged), suggesting that many may be aberrant promoters rather that repurposed from other cell types. Fisher's exact test odd ratio (OR) and *P*-values.

### Supplementary Fig. 7

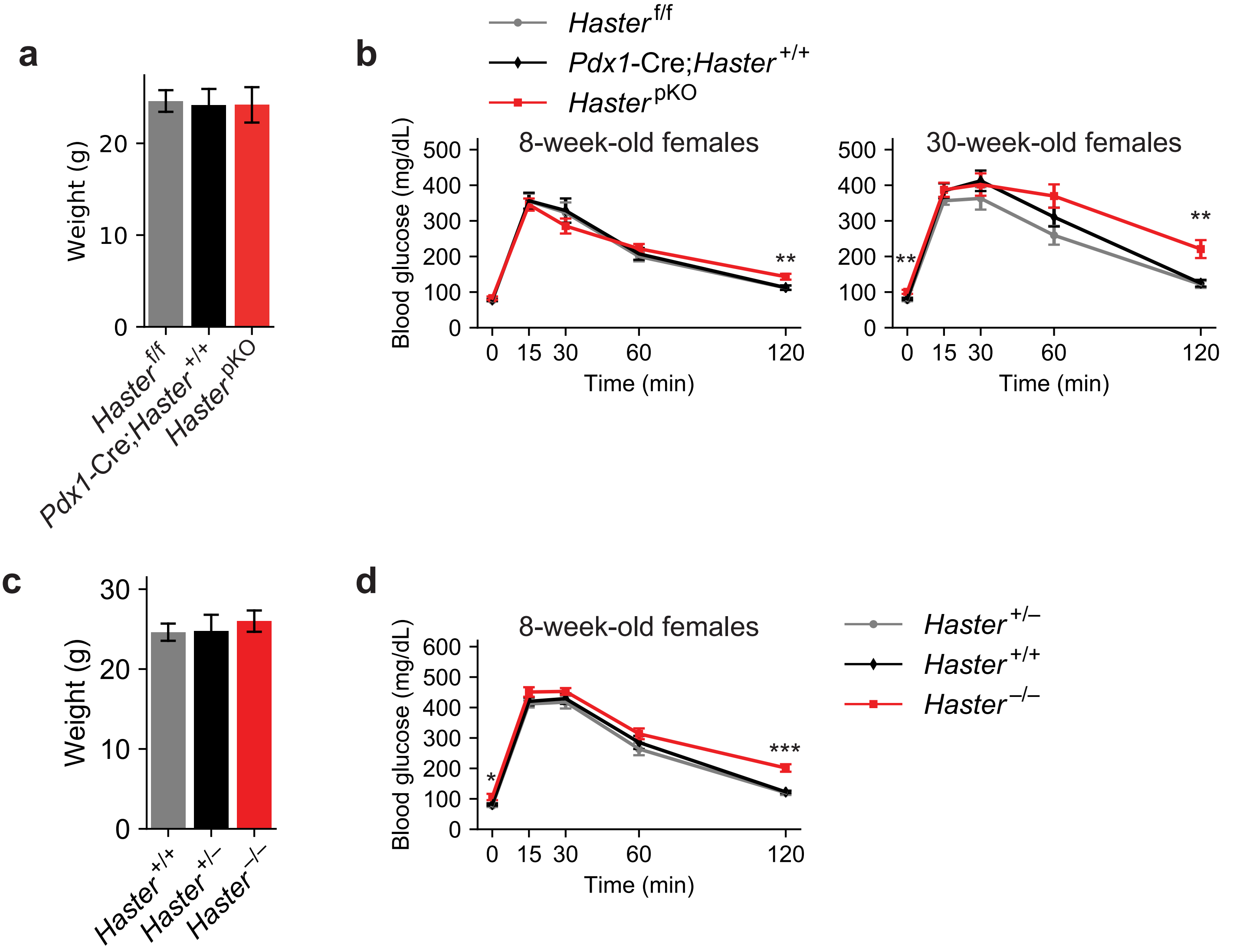


**Supplementary Fig. 7. Body weight of male *Haster* knockouts and glucose tolerance in female mice. a**, Body weight of males at 8 weeks of age. *Haster*$^{pKO}$ (n = 8), *Pdx1*-Cre;*Haster*$^{+/+}$ (n = 12) and *Haster*$^{f/f}$ (n = 8). Mean ± s.d.. b, Intraperitoneal glucose tolerance test in 8-week-old and 30-week-old female *Haster*$^{pKO}$ (n = 9), *Pdx1*-Cre;*Haster*$^{+/+}$ (n = 10) and *Haster*$^{f/f}$ (n = 10). Mean ± s.e.m., **P* ≤ 0.05 , ***P* ≤ 0.01 and ****P* ≤ 0.001 (two-tailed Student's t-test). **c**, Body weight of 8- to 10-week-old male *Haster*$^{-/-}$ (n = 9), *Haster*$^{+/-}$ (n = 12) and *Haster*$^{-/-}$ (n = 13). Mean ± s.d.. **d**, Intraperitoneal glucose tolerance test in 8- to 10-week-old females *Haster*$^{-/-}$ (n = 10), *Haster*$^{+/-}$ (n=10) and *Haster*$^{+/+}$ (n = 12). Mean ± s.e.m., **P* ≤ 0.05, ***P* ≤ 0.01 and ****P* ≤ 0.001 (two-tailed Student's t-test).

### Supplementary Fig. 8

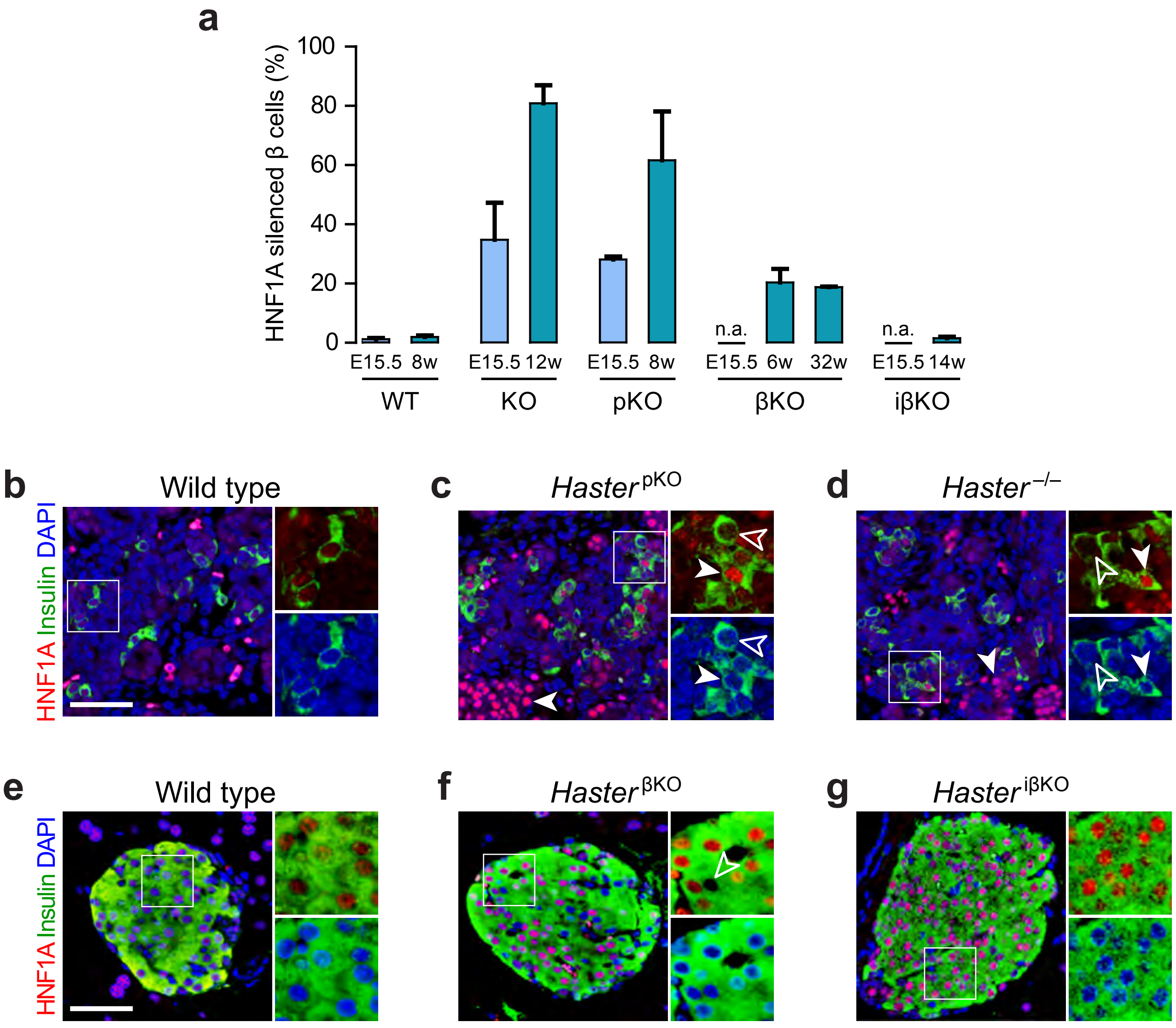


**Supplementary Fig. 8. Variegated HNF1A expression in *Haster* KO islet cells. a**, Relative quantification of HNF1A-negative β cells at the indicated age and genotype. Results show that HNF1A silencing correlates with time of *Haster* knockout, with higher silencing frequency after early deletion (*Haster* germline KO and *Haster* $^{pKO}$ models). HNF1A silencing increased with time in β cells from germline KO and *Haster* $^{pKO}$ models, but not when excision occurred in early β cells (βKO). No HNF1A silenced β cells were observed after *Pdx1*-Cre$^{ERTM}$-based tamoxifen-inducible excision in adult β cells (*Haster* $^{i\beta KO}$ model). n = 2 wild type embryos and n = 3 adult wild type mice; n = 2 *Haster* $^{-/-}$ embryos and adult mice; n = 3 *Haster* $^{pKO}$ embryos and adult mice; n = 2 *Haster* $^{\beta KO}$ 6- and 32-week-old mice; n = 2 *Haster* $^{i\beta KO}$ mice. Mean ± s.d.. **b-d**, Immunofluorescence for HNF1A and insulin in E15.5 (**b**) *Haster* $^{+/+}$, (**c**) *Haster* $^{pKO}$, and (**d**) *Haster* $^{-/-}$ pancreas. Solid arrows, insulin cells overexpressing HNF1A; hollow arrows, insulin cells lacking HNF1A. **e-g**, Immunofluorescence for HNF1A and insulin in adult (**e**) wild-type, (**f**) *Haster* $^{\beta KO}$ and (**g**) *Haster* $^{i\beta KO}$ pancreas. Arrows point to HNF1A-negative β cells. Most β cells from *Haster* $^{\beta KO}$ and *Haster* $^{i\beta KO}$ islets overexpress HNF1A. Scale bar, 50 μM.

#### Supplementary Fig. 9

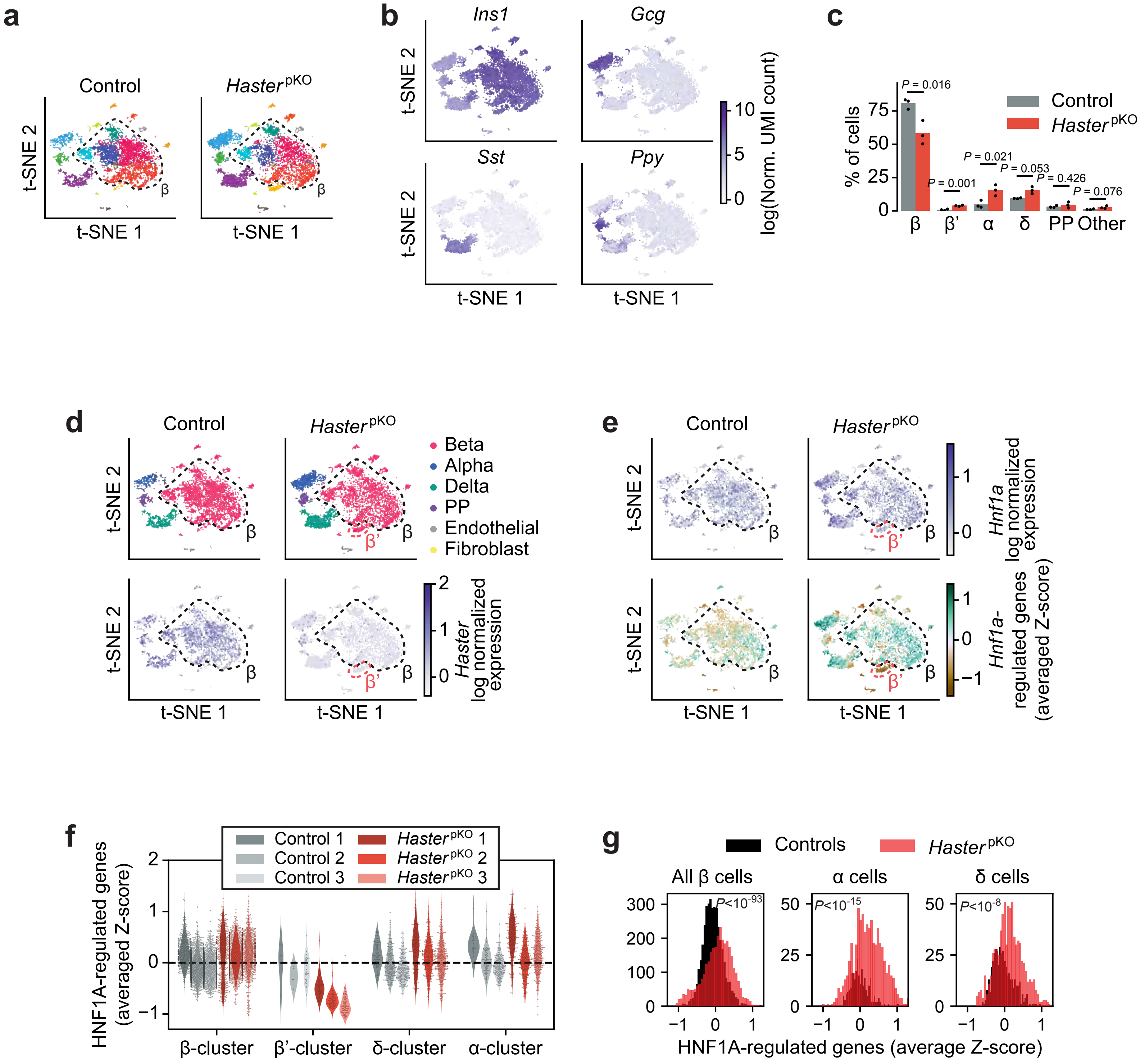


**Supplementary Fig. 9. Single-cell RNA-seq of *Haster*$^{pKO}$ islets. a**, scRNA-seq t-SNE plots of islet cells from female *Haster*$^{pKO}$ (4961 cells from triplicates) and controls (4646 cells from triplicates). A dotted line encompasses the main group of β cell clusters. **b**, t-SNE plots showing hormone expression. **c**, Relative proportions of islet cells. Each scRNA-seq cluster was assigned a cell type based on main hormone expression. β cell clusters were grouped, except for β' cell cluster that was only seen in *Haster*$^{pKO}$ cells. Mean ± s.d., two-tailed Student's t-test. Note that these differences in cell composition are not per se expected to cause diabetes. **d**, Haster mRNA (log normalized UMI count) in different cell types of control and *Haster*$^{pKO}$ islets. **e**, *Hnf1a* mRNA (log normalized UMI count) in top panels; and expression of Hnf1a-regulated genes (average Z-score of log normalized UMI count) in bottom panels. The black dashed line highlights the main β cell cluster, showing increased HNF1A-regulated genes in *Haster*$^{pKO}$, and the red dashed line highlights the β'cluster that is only observed in *Haster*$^{pKO}$ and shows decreased HNF1A-dependent gene expression. **f**, HNF1A-regulated gene expression (average Z-score) for different cell types in individual samples. **g**, Histograms showing the distribution of the HNF1A-regulated gene expression (average Z-score) for α, β and δ cells. Bins 40. The variance of HNF1A-regulated gene expression increased in *Haster*$^{pKO}$ α, β and δ cells, showing that HNF1A-regulated genes are either upregulated or downregulated in islet cells. Levene test *P*-values.

### Supplementary Fig. 10

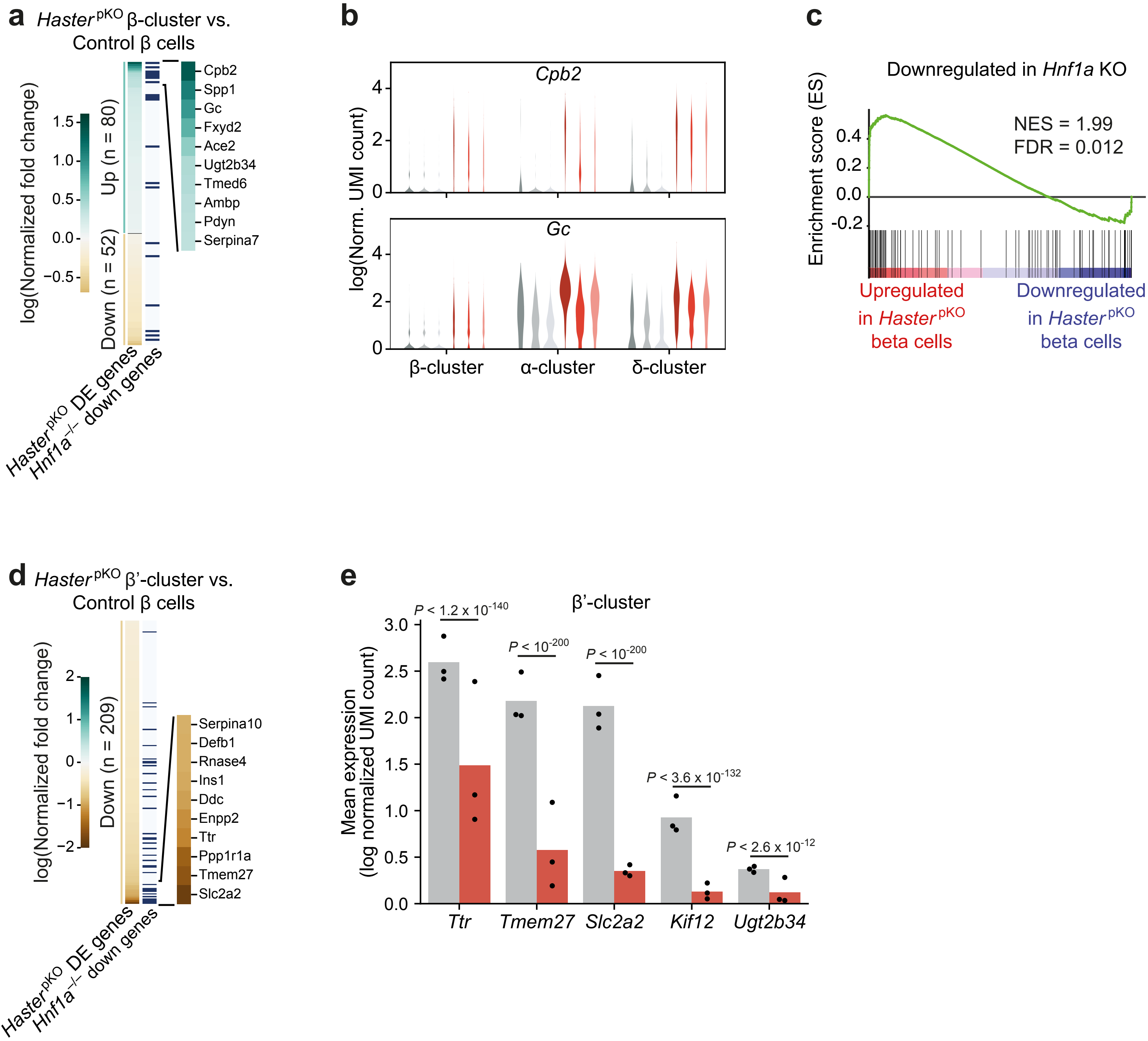


**Supplementary Fig. 10. Differential gene expression in *Haster* $^{pKO}$ β cells. a**, Genes differentially expressed in the major β-cell cluster of *Haster* $^{pKO}$ islets. Many of the most upregulated genes in *Haster* $^{pKO}$ islets are downregulated in *Hnf1a* $^{-/-}$ islets (blue horizontal lines). **b**, Examples of two genes that are known to be downregulated in *Hnf1a* $^{-/-}$ islets, *Cpb2* and *Gc*, and show increased expression in *Haster* $^{pKO}$ β cells. c, GSEA showing upregulation in *Haster* $^{pKO}$ β cells of genes downregulated in *Hnf1a* KO islets. **d**, Genes that are downregulated (combined $P \leq 0.05$) in *Haster* $^{pKO}$ β' cluster cells are often downregulated in *Hnf1a* $^{-/-}$ islets (blue horizontal lines). **e**, Expression of selected genes that are known to be downregulated in *Hnf1a* KO islets and are downregulated in *Haster* $^{pKO}$ β'cells. Dots are medians of samples (log normalized UMI count) and bars are means of 3 replicates. Wilcoxon Rank Sum test, *P*-values for the different biological replicates combined with Fisher's method.

### Supplementary Fig. 11

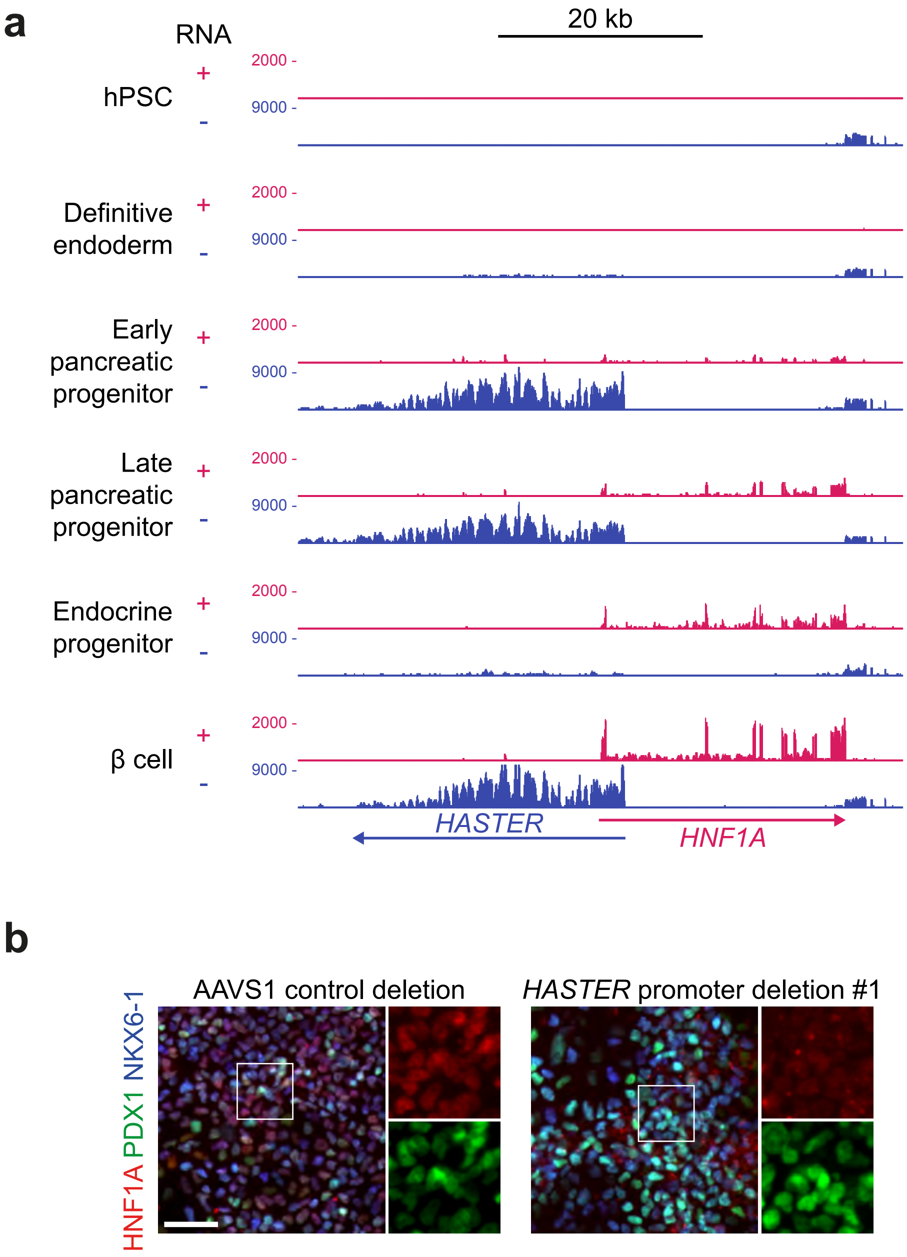


**Supplementary Fig. 11. *HASTER* deletion in human stem cell-derived pancreatic cells. a**, RNA-seq showing *HASTER* and *HNF1A* expression during hPSC differentiation to β cells (data from Alvarez Dominguez et al, Cell Stem Cell, 2020). **b**, Immunofluorescence for HNF1A and the pancreatic progenitor markers PDX1 and NKX6-1. Scale bar, 50 μM.

### Supplementary Fig. 12

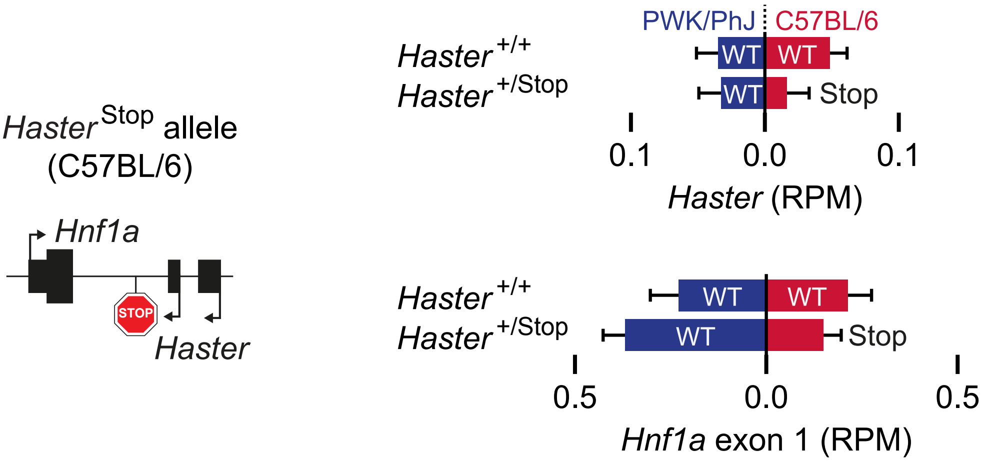


**Supplementary Fig. 12. *Haster* RNA or elongation are dispensable for the negative regulation of *Hnf1a* by *Haster*.** Strain-specific RNA expression from *Haster* $^{+/stop}$ C57BL/6;PWK/PhJ hybrid mice, showing that reducing *Haster* elongation in failed to increase *Hnf1a* expression from the same C57BL/6 allele.

### Supplementary Fig. 13

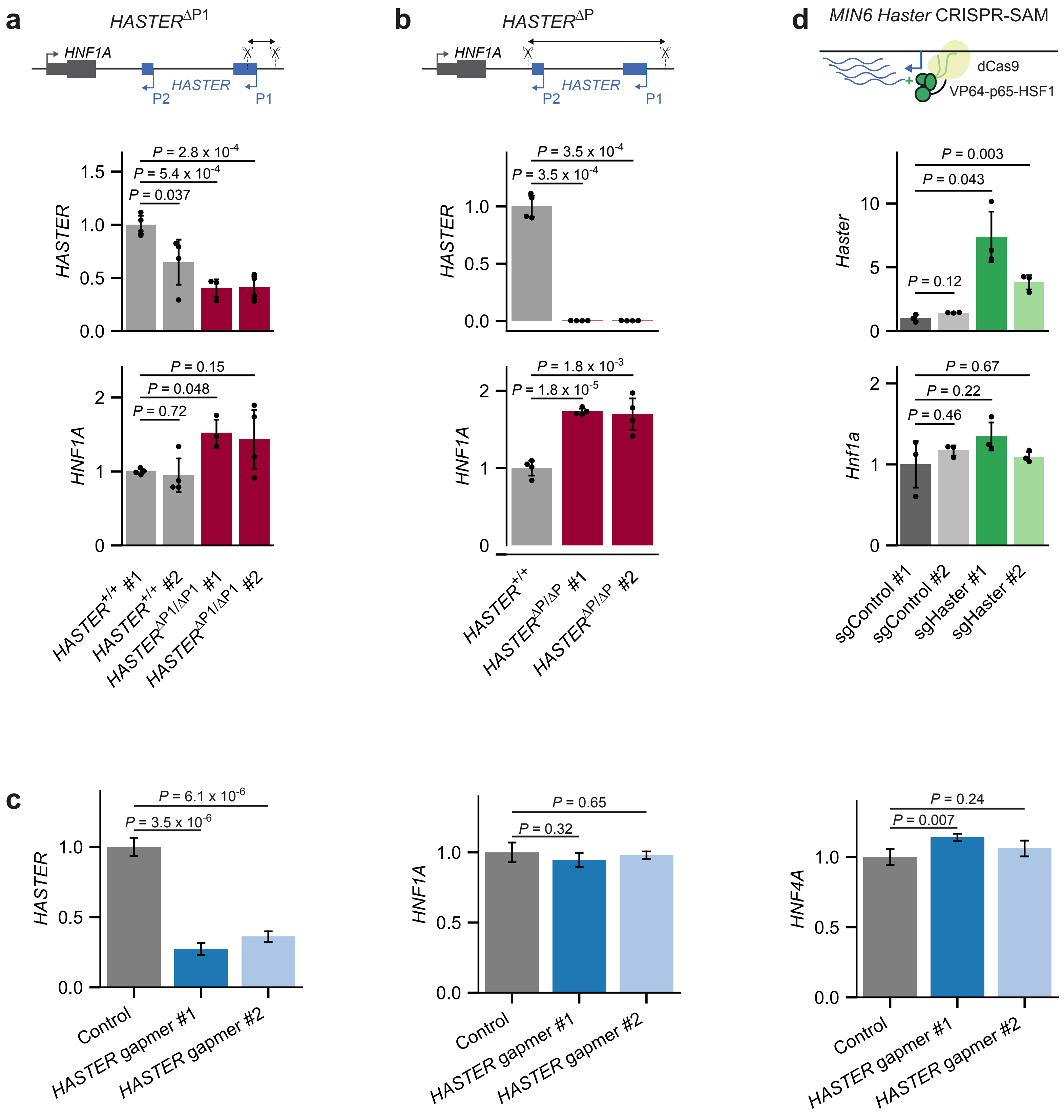


**Supplementary Fig. 13.** ***HASTER*** **perturbations in β cells. a**,**b**, *HASTER* and *HNF1A* RNA in EndoC-βH3 cells with clonal homozygous deletion of (**a**) *HASTER* P1 promoter (*HASTER*$^{\Delta P1/\Delta P1}$) or (**b**) *HASTER* P1 and P2 promoters (*HASTER*$^{\Delta P/\Delta P}$). Deletion #1 and #2 were generated with independent pairs of sgRNAs, *HASTER*$^{+/+}$ clones were transfected with sgRNAs targeting the AAVS1 locus. n = 4 clones per deletion. **c**, Two sets of locked nucleic acid oligonucleotides (GapmeRs) were used to elicit *HASTER* degradation in EndoC-βH3 cells, without significant changes in *HNF1A* or *HNF4A* mRNA. n = 3 nucleofections. **a-c**, Expression normalized by *TBP*. Mean ± s.d., two-tailed Student's t-test. **d**, *Haster* activation by CRISPR-SAM in MIN6 mouse β cells had no effect on *Hnf1a* expression. n = 3 lentiviral transductions. Expression normalized by *Tbp*. Mean ± s.d., two-tailed Student's t-test relative to the control #1 sgRNA.

### Supplementary Fig. 14

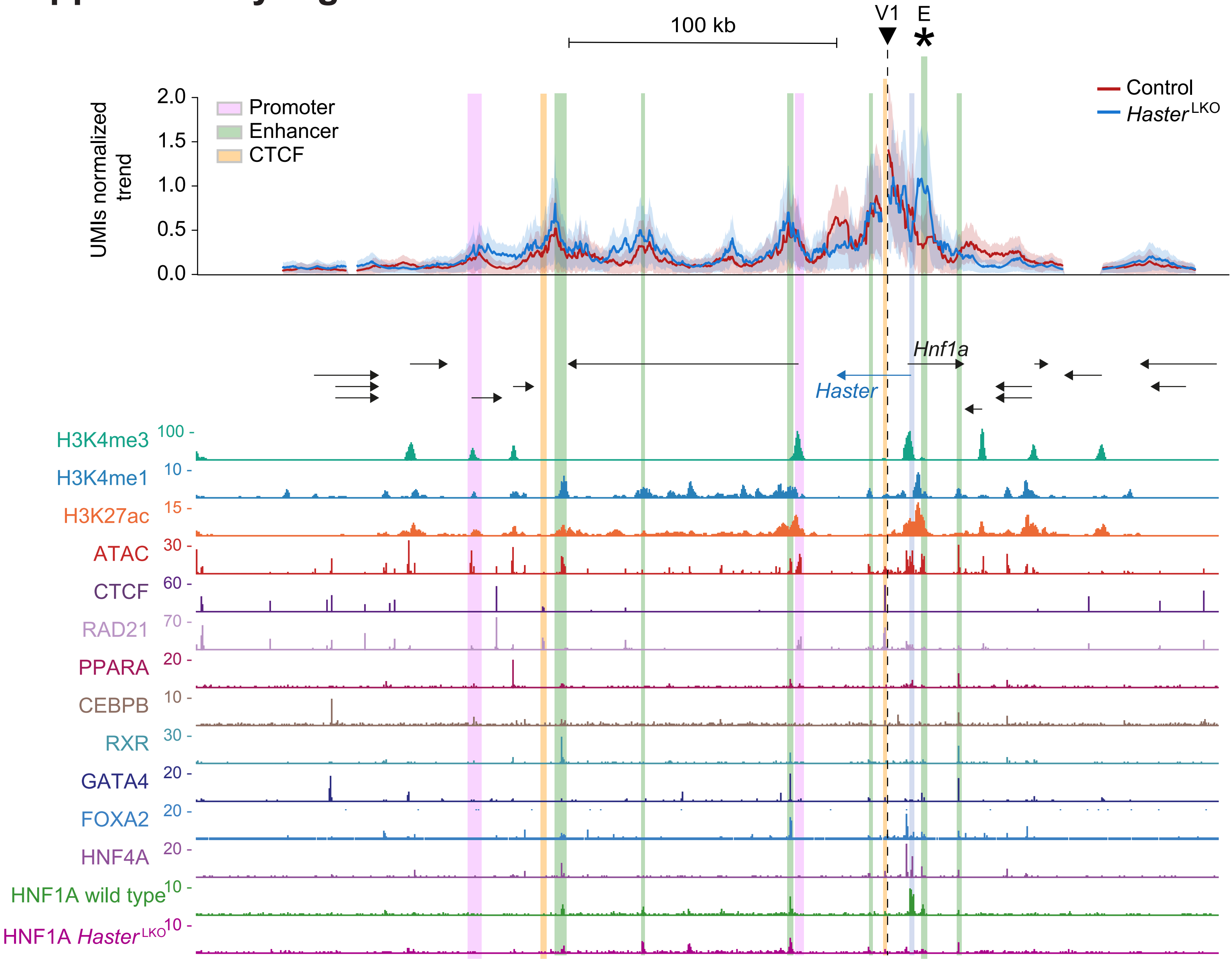


**Supplementary Fig. 14. Remodeling of local chromatin contacts in *Haster*$^{LKO}$ liver.** Top, UMI-4C profile trends using a viewpoint region upstream of *Hnf1a* (V1), near a CTCF-bound C site, in adult liver from n = 3 wild type (blue) or mutant (red) mice. Bottom, chromatin features and transcription factor binding in adult mouse liver. The *Hnf1a* upstream region contacts several enhancers, promoters and CTCF/cohesin sites in control and *Haster*$^{LKO}$ liver. The interaction between *Hnf1a* upstream region and E (asterisk) is increased in *Haster*$^{LKO}$ liver (see also **Fig. 7**). The region deleted in *Haster*$^{LKO}$ mice is highlighted in blue.

### Supplementary Fig. 15

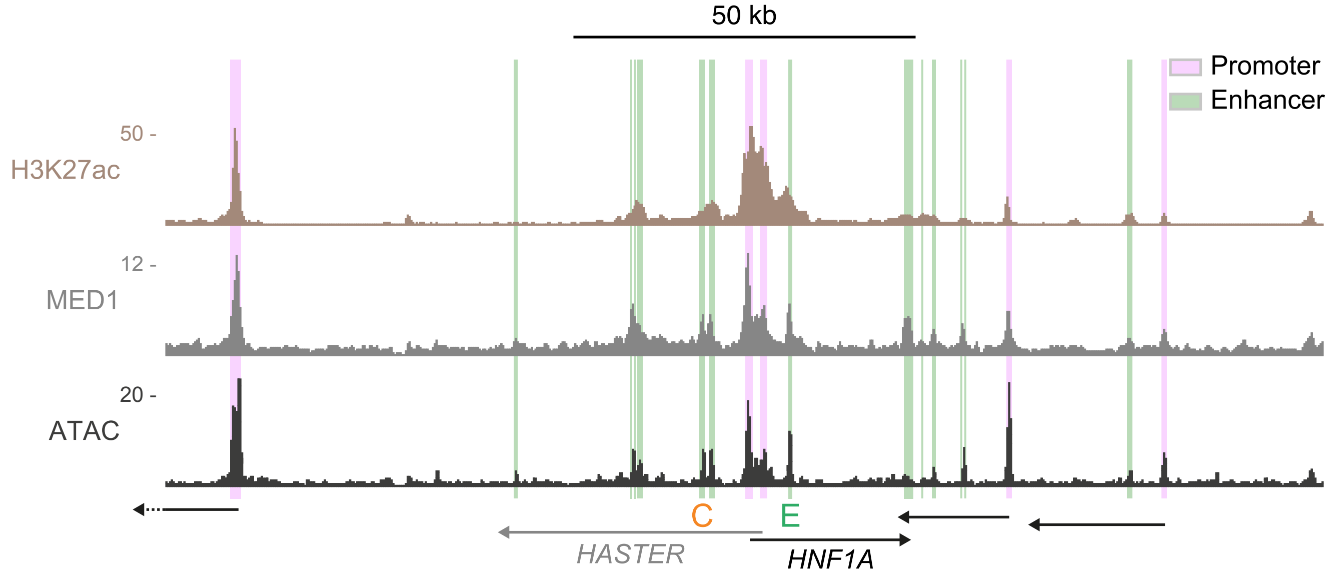


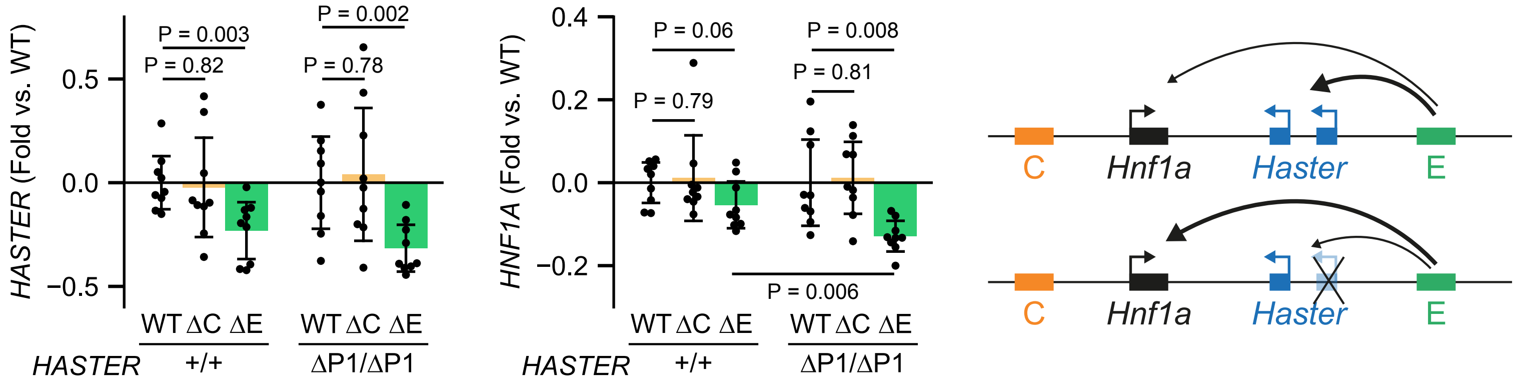


**Supplementary Fig. 15. E deletion partially prevents *HNF1A* mRNA increase in *HASTER* KO β cells. a**, Human islet chromatin marks showing the position of enhancers in the vicinity of *HNF1A*. **b**, *HASTER*$^{+/+}$ or *HASTER*$^{\Delta P1/\Delta P1}$ clone #1 cells carrying targeted deletions in C (ΔC), E (ΔE) or sgGFP as control (WT). *HASTER*$^{+/+}$ control and E deletion are identical to **Fig. 7e**. ΔC and ΔE were polyclonal deletions. Results are expressed as fold-differences relative to the parental *HASTER*$^{+/+}$ or *HASTER*$^{\Delta P1/\Delta P1}$ cells. This showed that ΔC has no effect on *HASTER* or *HNF1A*, ΔE had significant effects on *HASTER* but did not significantly affect *HNF1A* in wild type cells, yet showed a significant *HNF1A* reduction in *HASTER*$^{\Delta P1/\Delta P1}$ cells. This is shown in cartoon form in the right panel, whereby E predominantly enhances *HASTER* transcription, but enhances *HNF1A* in the absence of *HASTER*. Pool of n = 3 independent experiments with 3 pairs of sgRNAs for each deletion. *TBP*-normalized mean ± s.d.; two-tailed Student's t-test.
